# Effects of host DNA and k-mer size on metagenomic classification of historical samples

**DOI:** 10.1101/2025.04.05.647358

**Authors:** Siu-Kin Ng, Rafal M. Gutaker

**Affiliations:** Royal Botanic Gardens, Kew, Richmond, Surrey, United Kingdom

## Abstract

Microbiome composition and function are strongly influenced by environmental factors, with major shifts driven by intensified anthropogenic pressures over the past centuries. This timeframe extends beyond the scope of experimental and longitudinal studies traditionally used to investigate microbiome dynamics. Historical samples could provide important insights into the mechanistic consequences of anthropogenic pressures and the resulting shifts in microbial diversity and composition. Despite their vast potential, historical samples held in museums and herbaria worldwide remain underutilized for exploring host-microbiome interactions across broad temporal and spatial scales, owing to their incompatibilities with standard analytical pipelines and a limited understanding of optimal analytical parameters. Host DNA removal has conventionally been considered essential for taxonomic assignment of metagenomic reads, and may be particularly important when processing degraded DNA, but it is impractical for specimens whose host species lacks a reference genome. Here, using empirical and simulation datasets, we show that host DNA content has a negligible impact on microbial data analysis. Because DNA molecules from historical samples are highly fragmented and variable in length, we further analysed the effect of k-mer size on the classification of metagenomic reads from historical samples. To improve recall rate, we propose a simple two-step approach in which reads are classified against two databases constructed with long and short k-mer sizes. Using simulated and published datasets, we demonstrated that this approach outperforms single-step workflows in recovering microbial signals from reads across a wide range of lengths. Together, our findings provide a foundation for incorporating natural history collections into host-associated microbiome research, offering insights into the long-term effects of anthropogenic change on microbial communities.

## Introduction

The microbiome refers to a collection of microorganisms, which comprise bacteria, archaea, viruses, microbial eukaryotes, and the surrounding environmental conditions (Marchesi & Ravel, 2015). Emerging evidence suggests the importance of the resident microbiome in enhancing the fitness of host species, supporting adaptation to internal and external changes (Debray et al., 2023; Ley et al., 2008; Petersen et al., 2023; Xiao et al., 2022). For example, the soil microbiome contributes to plant growth through biogeochemical cycling of nutrients and organic matter (Banerjee & van der Heijden, 2023; Jansson & Hofmockel, 2020), the endophytic microbiome contributes to plant survival under drought (Naylor & Coleman-Derr, 2017; Trivedi et al., 2020) and the gut microbiome to the fitness and lifespan of Drosophila (Gould et al., 2018).

The field of microbial ecology aims to identify and characterize microbial community composition, function and their interactions with changing environments. Samples from controlled experiments (e.g. conventional case-control or longitudinal studies) are often sufficient to provide useful insights into the effects of short-term perturbations, such as dietary or seasonal climate shifts. The study of long-term disturbances or environmental changes (Gancz et al., 2023; Morais et al., 2024; Rout et al., 2023; Tiao et al., 2012), however, cannot be achieved with these samples. Historical samples, including those in herbarium or museum collections, have been preserved for centuries and offer a unique opportunity to examine the effects of long-term disturbance on hosts and their associated microbiomes using metagenomic approaches (Warinner et al., 2017; Weyrich et al., 2017). Examples of fundamental changes in the last two centuries include increased antimicrobial resistance in microbes in response to the introduction of antibiotics, and the decline of soil nitrogen fixing activity following the widespread application of synthetic fertilizers (Aeron et al., 2020; Naamala et al., 2016; Rout et al., 2024). Historical samples provide a global and extended temporal perspective that conventional studies simply cannot achieve (Burbano & Gutaker, 2023). They could be an important source to uncover the diversity of microbial community predating major anthropogenic activities.

Host DNA and low endogenous microbial biomass (Eisenhofer et al., 2019) make metagenomic analysis with historical samples challenging. Additionally, nucleic acid material preserved in historical samples is prone to long-term degradation and environmental contamination. The impact of host DNA content on taxonomic classification of microbial communities has been investigated (Pereira-Marques et al., 2019) and host DNA removal prior to data analysis has been seen as a necessary step. Several commercial DNA extraction kits are designed to perform differential lysis and digestion of host cells to deplete their DNA, however, these kits are only effective for human or animal host cells but not for plant nor host cells with complex cell wall structure (either fresh or historical). Alternatively, removal of host DNA can be performed *in silico* with the reference genomes of host species or closely related species. However, the breadth of microbe-host-environment interactions is best studied with diverse species including non-model organisms (e.g. diverse angiosperm plants (Zuntini et al., 2024)), where representative reference genomes are often absent.

Several critical questions regarding microbial analyses in historical samples remain unaddressed. First, is host DNA removal necessary when profiling the host-associated microbiome in historical samples, particularly those from non-model organisms? Although host DNA content may reduce the sensitivity for detecting rare taxa (Pereira-Marques et al., 2019), it remains unclear whether it significantly impacts microbiome analyses such as microbial diversity metrics. Second, the nucleic acids (either from host or microbes) preserved in historical samples are typically highly fragmented due to degradation processes such as cytosine deamination (Gutaker & Burbano, 2017). Previous studies have developed methods to recover very short DNA fragments (e.g. 25 bp) from historical samples (Glocke & Meyer, 2017; Harney et al., 2021) but it remains unclear how informative these short metagenomic reads might be. Without removing host DNA, short reads derived from the host may be misclassified as microbial or vice versa. However, discarding these short reads from downstream analyses is not ideal, particularly when working with precious museum or herbarium collections. Microbiome profiling often relies on alignment-free tools such as Kraken2 (Lu et al., 2022), where the sensitivity and specificity of classification are influenced by parameters such as k-mer size or seed length. By fine-tuning these parameters, short reads can still contribute to important insights. In this study, we explored the impact of host DNA on host-associated microbiome analyses using both simulations and published datasets from historical collections. We also evaluated parameters and strategies for analysing the metagenomic reads obtained from samples lacking host reference genomes.

## Materials and Methods

### Sampling, sequencing and collection of museum samples

A total of 864 metagenomes of contemporary and historical specimens were included in this study to investigate the impact of host DNA content on microbial analyses and to evaluate appropriate analytical strategies. For the contemporary specimens, 330 leaf metagenomes (PRJEB45634) were generated from 110 rice cultivars (genotypes) that were arranged in a randomized complete block design (three plots for each cultivar) (Su et al., 2022). For the historical samples, 90 Asian rice (PRJNA1282210) (Alam et al., 2025), 37 common ragweed (PRJEB48563) (Bieker et al., 2022), and 402 bumblebee (PRJEB52125) (Mullin et al., 2023) specimens were included. In addition, five phyllosphere metagenomes from herbarium rice species were generated in this study, and the raw sequencing data were deposited in the EBI nucleotide archive (PRJEB87646). Raw sequencing data for the public datasets were retrieved from the NCBI Sequence Read Archive. Genome assemblies for the host species were downloaded from NCBI GenBank or EBI ENA: rice (*Oryza sativa* AGIS1.0; GCA_034140825.1 for reference genome; MZ424305.1 for chloroplast), common ragweed (*Ambrosia artemisiifolia*; GCA_024762085.1 for reference genome; MG019037.1 for chloroplast) and bumblebee (*Bombus lapidarius; Bombus_lapidarius_EIv1*; PRJEB51891). The datasets and genomic sequences included in this study are summarized in Supplementary Table 1.

In the k-mer analysis and simulation study, we considered the soil microbiome as a proxy for the host-associated microbiome because of the high microbial diversity in the soil environment and its biological relevance to the plant species included in this study. For the k-mer analysis, a microbial k-mer profile was created from 107 pan-genomes of soil microbes published along with the catalogue of soil metagenome-assembled genomes (SMAGs; PRJNA983538). The SMAGs catalogue contains 21,077 genomic bins reconstructed from 3,304 soil metagenomes and resolved at the species level (Ma et al., 2023). These pan-genomes were generated from 2,200 species-level genomic bins (a subset of 21,077 genomic bins) with >10 high-quality metagenome-assembled genomes (MAGs) by clustering conspecific protein sequences at 90% amino acid identity. They also represent unique lineages of soil microbes and therefore provide a broader spectrum of unique sequences commonly found in the soil microbiome, serving as a “sequence/k-mer pool” of soil microbes. We compared k-mers in these microbial pan-genomes to those from host species to assess how likely microbial and host k-mers are matched simply by chance. Wherever possible, curated reference genomes (sequence accession with a GCF_* or GCA_* prefix) were used.

For the simulation study, we shortlisted 120 bacterial species based on the number of species per genus (Supplementary Table 2) in the SoilGenomeDB database (Edwin et al., 2024) and completeness of their genomes. This allowed read simulators to synthesize reads evenly across complete genomes or fragmented genomic sequences (contigs). In addition, each simulated dataset (6,000 in total) contained 1 million reads (resembling the metagenomes obtained from historical samples). Inclusion of too many microbial genomes in the simulation would end up with most of the microbial taxa failing the downstream filtering steps due to a low number of reads assigned for each taxon. Therefore, the use of 120 species was computationally and analytically practical for the simulation study without deviating substantially from empirical expectations. Microbial reference sequences were downloaded from NCBI GenBank using the “download” command of the NCBI Datasets command-line tools.

### Collection and processing of herbarium rice specimens

Five leaf metagenomes were characterised from rice species collected between 1906 and 1914 in the Philippines and stored in the herbarium collection of the Smithsonian Institution. Briefly, a small fragment of leaf (approximately 20 mm x 5 mm) was removed and processed in the semi-clean room facilities (without air filtering) at the New York University Center for Genomics and Systems Biology. Standard precautions for handling historical and ancient DNA were taken (disposable personal protective equipment, use of dedicated polymerase chain reaction (PCR) UV cabinets without air circulation, frequent cleaning regimes and UV-sterilization). Sampled tissue was ground ‘wet’ with N-Phenacylthiazolium bromide (PTB) lysis buffer at 37°C. Subsequently, DNA isolation and genomic library preparation protocols (without uracil-DNA glycosylase treatment) for historical plant tissues were adopted from our previous work (Latorre et al., 2020) with the following modifications: indexing PCR was carried out with Agilent Pfu Cx polymerase. Libraries were amplified with 10 PCR cycles, quantified using real-time quantitative PCR, normalized and pooled to equimolar concentrations. Libraries were then sent to Macrogen for sequencing on a NovaSeq 6000 S4 flow cell.

### Preprocessing of metagenomic data

Raw demultiplexed reads were processed with the nf-core/mag pipeline, an open-source and community-curated best-practice bioinformatics pipeline for preprocessing, assembly, binning and annotation of metagenomes based on the Nextflow nf-core framework (Ewels et al., 2020; Krakau et al., 2022). Quality control of the raw metagenomic sequencing reads in FASTQ format was performed using FastQC v0.12.0 (Andrews, 2015). Reads were quality filtered and trimmed with the all-in-one fastp preprocessing tool using default settings (Chen et al., 2018), followed by merging using AdapterRemoval v2.3.4 (Schubert et al., 2016). Reads mapped to the PhiX sequence using Bowtie2 v2.3.4 were discarded (Langmead & Salzberg, 2012).

### Host DNA removal and taxonomic classification

The impact of host DNA content on metagenomic analyses was evaluated by comparing the microbial community metrics in the same samples processed (i) with host DNA removal or (ii) without host DNA removal (Figure 1). Host DNA removal was carried out by aligning filtered reads to the host genome using Bowtie2 with default settings. Reads that were not mapped to the host genome were then taxonomically annotated. Taxonomic classification of metagenomic reads was performed using Kraken2 v2.0.9 (Wood et al., 2019) with a confidence value of 0.4 against the PlusPFP database (prebuilt with k=35, version 2024.06.05), which contains sequences of archaea, bacteria, virus, plasmid and human, as well as protozoa, fungi and plants. The selected confidence value could achieve a reasonable classification rate without introducing a high false positive rate (Supplementary Figure 1). Kraken2 classifies metagenomic reads by mapping all k-mers to the lowest common ancestor (LCA) of all genomes present in the database. Non-microbial taxa (e.g. those assigned to eukaryotic species or viruses) were excluded from further analysis. Taxa were further filtered using a minimum read count of 50 (100 for contemporary rice samples due to their higher average read number) and a prevalence cutoff of 2% of samples (it was determined empirically so that taxa to be detected in at least 2 samples; this prevalence cutoff was practically disabled for small datasets: herbarium ragweed N=37 and rice N=95). Kraken2 was used to profile metagenomic reads in this study as it can achieve a better overall performance than other metagenomic profiling tools (Wright et al., 2023).

**Figure 1.**
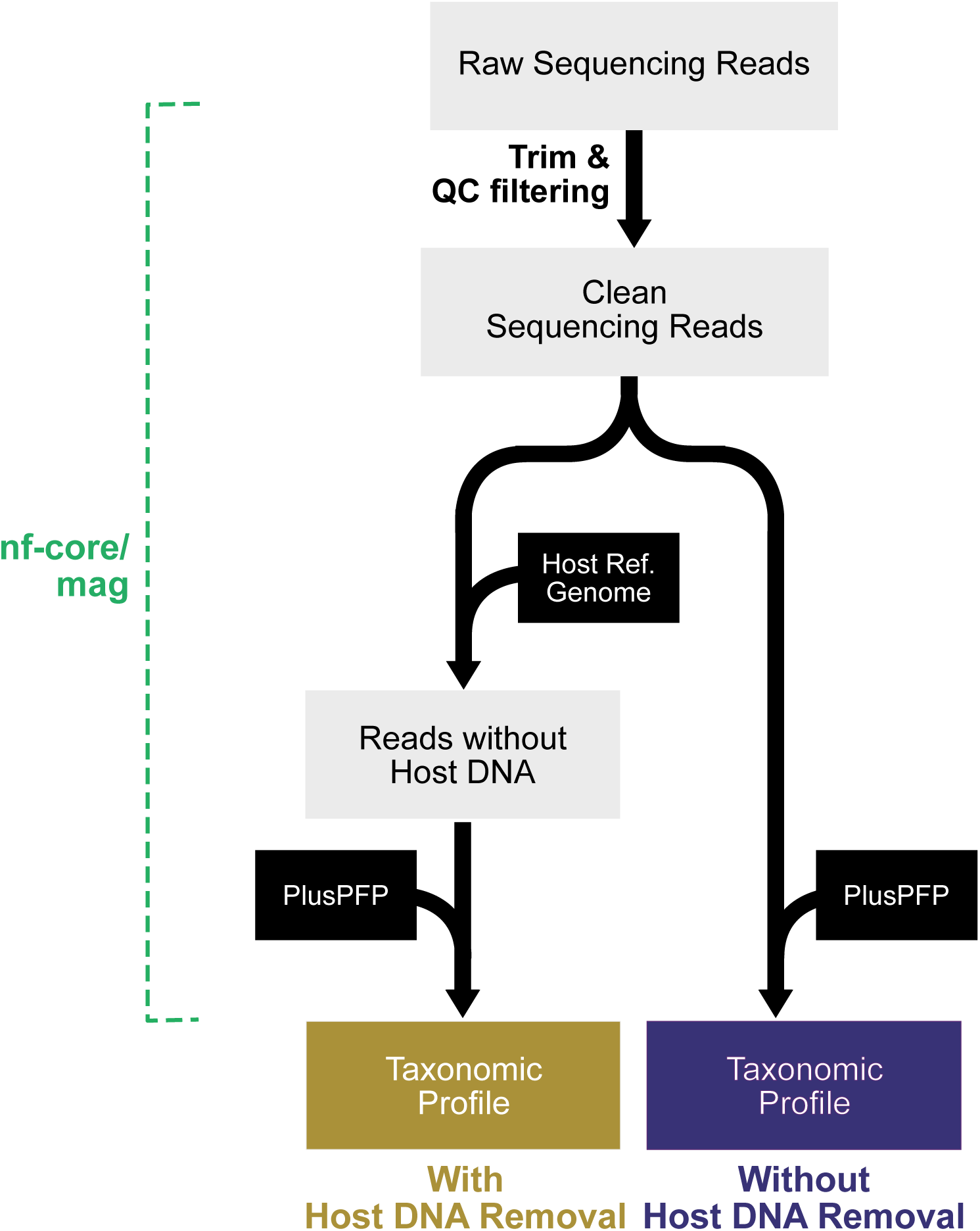
The bioinformatic workflow. The Nextflow metagenomic analysis workflow (nf-core/mag) was used in this study. Two separate workflows (gold for the workflow with host DNA removal and purple for the workflow without host DNA removal) were performed in parallel with identical steps except for the host removal step prior to taxonomic classification.

### K-mer analysis of host and associated microbiome

Misclassification of host DNA as microbial (cross-domain) may be a concern if host DNA is not removed. Estimating the host genomic sequences that are identical to microbial genomes (shared k-mers) could provide a theoretical assessment of how likely host sequences are to be misclassified as microbial purely by chance, based on the ratio of shared to distinct k-mers between host and microbes. Briefly, genomic sequences of host and microbes were first decomposed into constituent k-mers. For rice and ragweed, the reference genomes listed above were used to generate their k-mer profiles. The k-mer profile of the soil microbiome was generated collectively from 107 pan-genomes of soil microbes. Shared k-mers occur in the genomes of both host and microbes, whereas distinct k-mers occur in only one of them. Overlap between host and microbes was measured using Jaccard index, 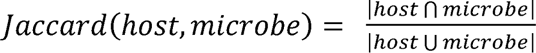. Counting of k-mers shared between host and microbial genome sequences was carried out with the “count” command from Jellyfish v1.1 (Marcais & Kingsford, 2011) with five different k-mer sizes at intervals of 3-4 nucleotides (11, 15, 18, 21 and 24). K-mer sizes larger than 24 was not evaluated as the analysis of large k-mer values using Jellyfish became computationally intensive. Also, we observed that the number of shared k-mers decreased substantially when k-mer size became larger than 21. K-mer substrings were exported to FASTA format with the Jellyfish “dump” command. Sequence complexity of k-mers was estimated as the Wootton-Federhen complexity score using the “sequence_complexity” function of the R package universalmotif v1.3 (Tremblay, 2024). The value of k-mers was used as window size, e.g. window.size=15 for k=15.

### Simulation of metagenomic datasets for historical samples

No *a priori* information on true microbial content in the contemporary or historical samples is available. To evaluate the effect of search parameters such as k-mer values, complexity of host genome and their DNA content in a sample, we performed a simulation study with datasets synthesized using gargammel v1.1.4, a new generation sequencing read simulator for ancient DNA (aDNA) (Renaud et al., 2017). The simulated dataset comprises three components: endogenous (***endo***), bacterial (***bact***) and contaminant (***cont***) DNA. We considered three plant host species to evaluate the impact of host genome size and complexity: rice (*Oryza sativa* AGIS1.0), maize (*Zea mays*; GCF_902167145) and bread wheat (*Triticum aestivum*; Genome assembly IWGSC CS RefSeq v2.1; chromosomes only). For the soil microbes, the reference genomes of 120 shortlisted bacterial species were used. Human genome (GRCh38.p14), *Cutibacterium acnes* (strain NBRC 107605; NZ_AP019723) and *Staphylococcus epidermidis* (strain ATCC 14990; NZ_CP035288) were included as contemporary contaminants. For each plant species, we created 100 input sequence sets of soil microbes with randomly assigned abundance. For each input sequence set, 20 simulated datasets consisting of 1,000,000 paired-end reads with a read length of 75 bp (HiSeq 2500) were iteratively generated from varying proportions of input sequence components ***bact*** (0.01, 0.05, 0.1, 0.3 and 0.45), ***cont*** (0.01, 0.05, 0.3 and 0.45) and ***endo*** (1.0 – (***bact*** + ***cont***)). To incorporate DNA fragmentation, consistent with the historical nature of DNA in museum samples, into the simulated datasets, the length profile of merged reads in a herbarium rice sample (HRB0263L) was created by tallying the length of merged reads using readlength.sh from the BBMap package (Bushnell, 2014) with the settings (bin=1 and n=1,000,000). The settings for damage patterns (nick frequency *v*=0.03; length of overhanging ends *l*=0.4; probability of deamination of cytosines (Cs) in double-stranded parts *d*=0.01; and probability of deamination of Cs in single-stranded parts *s*=0.3) were adopted from the DNA isolated from Pleistocene organisms (Briggs et al., 2007). These settings were used to recreate the damage patterns typical of samples that are 10,000 year old or older. We adopted them to generate datasets representing a worst-case scenario. In total, 6,000 simulated datasets were generated across three plant species.

The effect of k-mer values on taxonomic classification was assessed with databases built with different k-mer values. A total of 79,004 unique reference sequences (derived from prokaryotes, virus, plasmid and human) that were included in the Kraken2 Standard database (version 2/26/2026) were retrieved from NCBI. The sequences were used to build reference databases with k-mer sizes of 21, 24, 28, 31 and 35 using the kraken2-build command. The k-mer values of 11, 15 and 18 were found to be not informative in the previous section and therefore not evaluated in this analysis. The simulated datasets were processed and analysed with the two workflows as described in the previous section. The misclassification rate of host DNA was determined by taxonomy ID assigned to each read based on the output of classified reads exported from Kraken2. Any host-derived reads classified into any prokaryotic clades were considered as misclassified. Reads with a taxonomy ID at the species level or below (e.g. subspecies or strain) were considered species-level resolved.

### Statistical analysis

All statistical methods and visualization were performed in R v.4.2.2 with the corresponding R-packages (phyloseq v1.52, vegan v2.7, ggplot2). Before diversity analyses, samples were normalized by rarefying (down-sampling) their sequencing depth to the minimum value of bacterial read sums in their respective dataset (contemporary rice=60,713; herbarium rice=5,099; herbarium ragweed=7,306; museum bumblebee=7,006). Rarefaction analysis was performed by subsampling read counts from the taxon table using the rarefy_even_depth function from the phyloseq package. Samples with fewer than 5,000 reads were excluded from the diversity analysis. Alpha (within-sample) diversity (Chao1 and Shannon index) was calculated using the estimateR and diversity functions from the vegan package to provide an overview of the richness and diversity of microbial communities. Pairwise comparisons between the two pipelines with and without host genome removal were assessed with the paired Wilcoxon signed-rank test with continuity correction. All *p*-values were adjusted with the conservative Holm-Bonferroni method. Beta diversity (between samples) was evaluated by the analysis of similarities (ANOSIM) to test the difference in the composition of microbial communities using the anosim function (999 permutations) from the vegan package. Bray-Curtis distance metrics at the species level were used to perform two-dimensional principal coordinate analysis (PCoA) plots to visualize the differences among microbial communities. Taxa with a read count less than 10 were considered absent. The precision and recall of taxonomic classification for each read in the simulation datasets were evaluated by comparing the assigned taxonomy IDs with the true taxonomy IDs of the microbial species from which the reads were generated. Only species level assignments were considered; reads assigned to higher taxonomic ranks (e.g. genus level) were treated as false negative (FN). Reads whose assigned taxonomy IDs did not match the true taxonomy IDs were classified as false positive (FP), while only those with correctly matched taxonomy ID were considered as true positive (TP). Precision was calculated as TP / (TP + FP), and recall as TP / (TP + FN). For the two-step classification approach, reads were treated as FP if they were assigned an incorrect (species-level) taxonomy ID during the first stage. Reads were considered TP only if they received the correct taxonomy ID in the first stage, or if they were initially unassigned (FN) but correctly classified during the second stage. Some of the analyses in this study were conducted on CropDiversity-HPC (Percival-Alwyn, 2024).

## Results

### Negligible effect of host DNA on taxonomic assignment of metagenomic reads

In this study, we compared the workflows with or without host DNA removal to evaluate their impact on taxonomic profiling of metagenomic reads from contemporary and historical specimens of three host species. During the host DNA removal step, reads were discarded if they were aligned to the reference genomes of their respective host. Overall, these metagenomic datasets covered a wide spectrum of host DNA content (including the reads mapped to non-prokaryotic species), ranging from less than 0.1% to over 80% (Supplementary Table 3 and Supplementary Figure 2). Metagenomic reads were then taxonomically annotated with Kraken2 using the prebuilt PlusPFP database (Figure 1). The rice reference genome was included in the database, whereas the closest available species of the common ragweed was the common sunflower (*Helianthus annuus*), which is in the same Asteroideae subfamily. The database contained no comparable insect genomes for the bumblebee dataset (Supplementary Table 4). Together, this represents three scenarios of reference genome availability when annotating metagenomic reads from non-model organisms. In all scenarios, host DNA detected in downstream analyses was markedly reduced when host DNA removal was performed (Supplementary Figure 2). On average, the number of host-derived reads (from either plant or metazoa) differed by about a hundred-fold between the workflows (Supplementary Table 3). In contrast, the numbers of reads assigned to bacterial taxa were stable between workflows (∼1.0-fold), regardless of whether host DNA was removed (Figure 2, Supplementary Tables 3 and 5).

**Figure 2.**
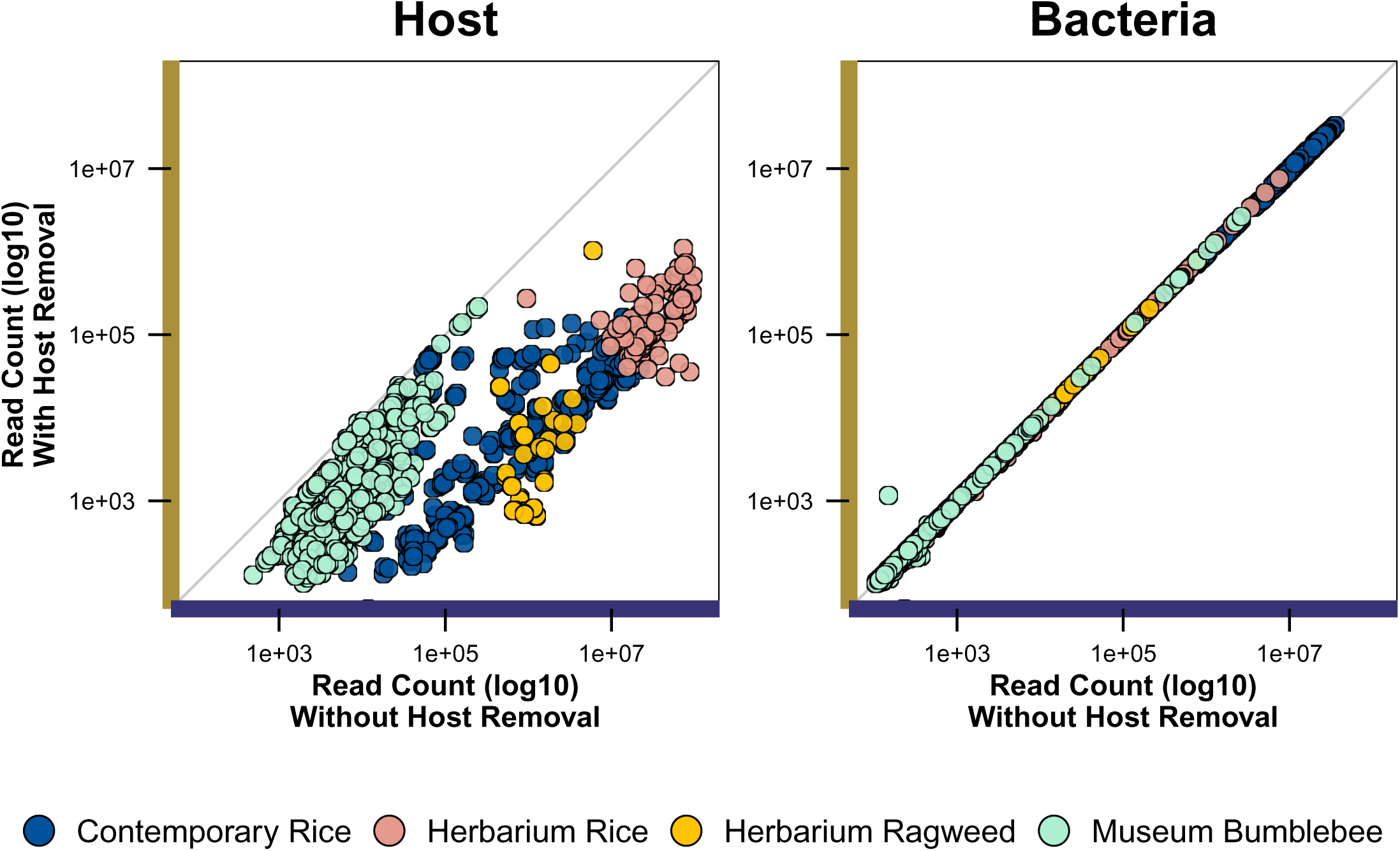
The number of sequencing reads (log10) assigned to host (left panel; plant or metazoa) and microbial taxa (right panel) with (y-axis; gold) and without (x-axis; purple) host DNA removal. Each circle represents a sample and is color-coded by their host species. Contemporary rice samples (blue circles) and historical rice samples (pink circles) were displayed separately despite the same host rice reference being used.

Next, we examined the taxonomic assignments in greater detail. To minimise false positives, prevalence (present in ≥2% of samples) and abundance (a minimum read count of 100 for contemporary rice and 50 for all other datasets) filters were applied to all detected taxa. Before filtering, the numbers of taxa ranged from 4,000 to 9,000, regardless of host DNA removal (Table 1). Most of these taxa (91-97%) had a relative abundance of 0.01% or lower and were removed as false positives in subsequent filtering steps (Supplementary Figure 3). For the contemporary rice samples, the same number of taxa (390) were retained from both workflows (with/without host removal) after filtering. Of these 390 taxa, 55 (14.1%) were highly variable (more than 10% difference) in their assigned read counts between the workflows, but their average read counts were lower than those of the taxa that were consistent between the workflows (Supplementary Figure 3). For the historical samples, a large portion of the detected taxa (>90%) were filtered out. After filtering, the museum bumblebee samples had the fewest taxa (113), whereas 853 taxa were retained in herbarium rice samples. The inclusion of the host DNA removal step changed the number of filtered taxa by only 0.5%, and almost all of the affected taxa were low in abundance (Table 1 and Supplementary Figure 3). No marked differences in microbial composition at the phylum or genus level were observed between the workflows in historical samples (Supplementary Figure 4). Overall, false-positives and low-abundance taxa were sensitive to taxon filtering but not to host DNA removal.

**Table 1.** Summary of the number of detected taxa (at the species level) between datasets with and without host DNA removal.

### No substantial change in metrics of microbial diversity even without host DNA removal

Without host DNA removal, misclassification of host reads as microbial (cross-domain) may be a concern, potentially leading to false detection of microbial taxa that are not present in the samples and distorting diversity metrics. We therefore assessed whether a host DNA removal step shift richness metrics and community diversity by measuring alpha and beta diversity, respectively. Rarefaction analysis indicated no noticeable difference in the number of detected taxa between workflows (Supplementary Figure 5). Alpha diversity was evaluated with two metrics: Chao1, a species richness estimator that accounts for rare or undetected species, and the Shannon diversity index, which combines richness and evenness. For the historical samples (Figure 3b-d), Chao1 values did not differ significantly (*p*>0.5) between the workflows with and without host DNA removal, with mean values of 59.3 ± 23.9 and 57.7 ± 23.4, respectively. The Shannon diversity index showed a similar pattern (*p*=0.43; 1.18 ± 0.72 with host removal and 1.17 ± 0.72 without; Figure 3a). For the contemporary rice samples, microbial diversity differed significantly between workflows because of a slight increase in species richness in the workflow without host DNA removal, but the effect size was very small. On average, an additional 0.67 taxa (median=1) were detected when host DNA was not removed, and the read counts of these additional taxa were only marginally above the abundance filter threshold (a minimal read count of 100). The effect size, measured by Cohen’s d, was very small (d=0.0341), indicating that the difference was statistically significant but practically negligible. Community composition, as measured by ANOSIM, also showed no detectable difference between the two workflows across all datasets (Figure 3).

**Figure 3.**
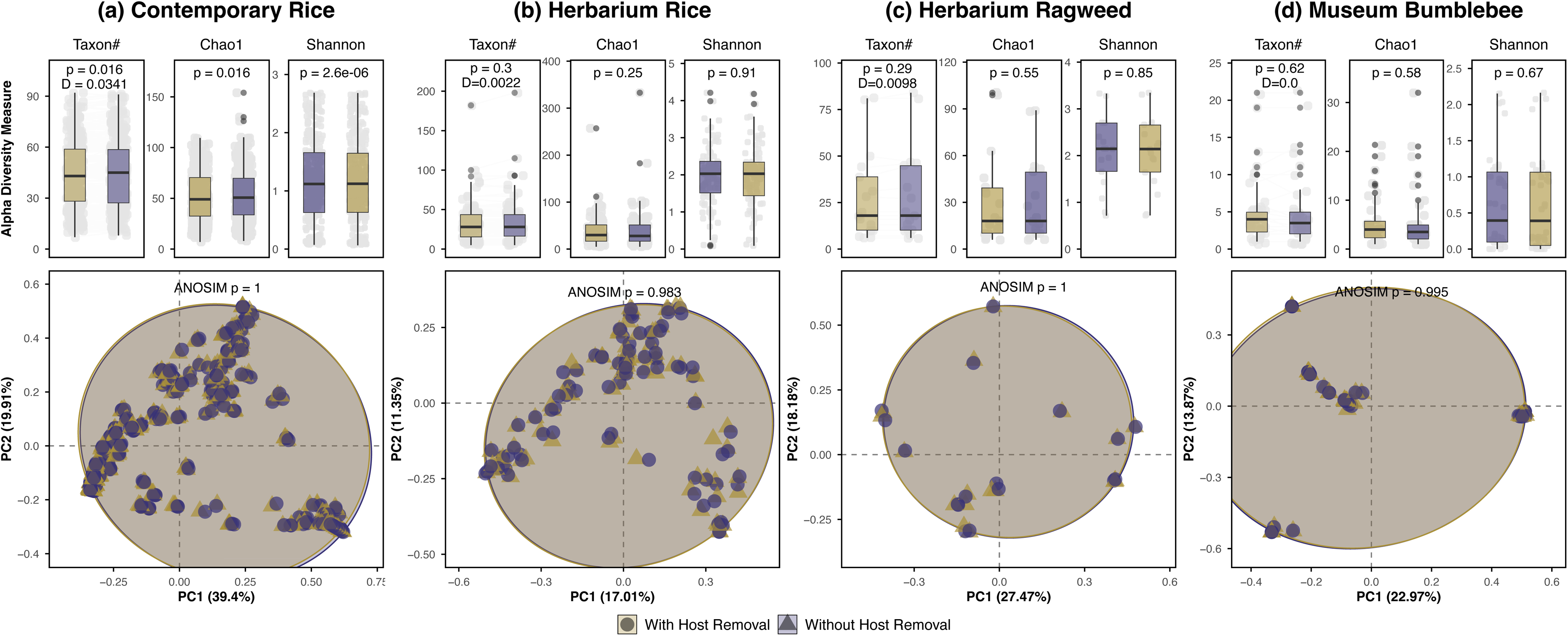
Alpha and beta diversity of the microbial community in the samples of (a) contemporary rice, (b) herbarium rice, (c) herbarium common ragweed and (d) museum bumblebee between the workflow with (gold) and without (purple) host DNA removal. Taxon richness and diversity are shown as the number of observed taxa, Chao1 and Shannon diversity indices (top panels) based on the mean values of 100 datasets rarefied to the read depth of 10,000. The structure of the microbial community obtained (bottom panels) from both workflows (gold triangles for the dataset with host removal; purple circles for the same dataset without host removal) was compared using principal coordinate analysis (PCoA) based on the Bray-Curtis dissimilarity index. *P*-value for microbial diversity was obtained from the Wilcoxon signed-rank test. The *p*-value of analysis of similarities (ANOSIM) was based on 999 permutations. The effect size for the number of observed taxa is shown in Cohen’s d.

### Optimal settings for classifying metagenomic reads from historical samples

K-mer analysis was performed on two plant species and a soil microbiome to assess the potential misclassification of host sequences as microbial. Unsurprisingly, the number of k-mers was strongly correlated with genome size and k-mer size (Supplementary Table 6). The number of distinct k-mers in the rice genome increased from 2,097,152 at k=11 to 282,788,287 at k=24, rising substantially between k=11 and k=15 and stabilizing from k=21 onward. A similar pattern was observed in the ragweed genome. Because the ragweed genome (1.3 Gb) is about three times the size of the rice genome (391 Mb), the number of distinct k-mers at k=24 was correspondingly 2.3-fold larger in ragweed. In the soil microbiome, the number of distinct k-mers was determined by species richness, which in this analysis was reflected by the total size of all pan-genomes (388 Mb), roughly equal to the size of the rice genome. Nonetheless, approximately 33% more distinct k-mers (0.37 billion) were obtained at k=24 than from rice (0.28 billion).

The number of shared k-mers between the host and the soil microbiome was measured using the Jaccard index, a metric of similarity based on the number of elements shared between two sets. At k=11, the host and the soil microbiome shared all k-mers, as reflected by a high Jaccard index (Figure 4a-4b; Supplementary Figure 6; Supplementary Table 6). The proportion of shared k-mers decreased gradually as k-mer size increased, with the Jaccard index dropping to 0.01 at k=18 and above (Figure 4b). The occurrence of k-mers across the host genomes was not uniform: some k-mers occurred only once, whereas others occurred more than a thousand times. We stratified k-mers by their occurrence frequency in the host genomes and reasoned that, if matches between host and microbial k-mers arose purely by chance, the frequency distribution of shared k-mers would closely follow that of the host genome (the expected distribution). The distribution of shared k-mers was largely consistent with the expected distribution at smaller k-mer sizes (k=18 or smaller) but deviated from it at larger sizes (Figure 4c). At k=24, the percentage of high-occurrence k-mers (5,000 or more) was markedly higher than expected. Furthermore, the sequence complexity of shared k-mers was significantly higher than that of the all k-mers when k-mer size became larger than 15 (Figure 4d), indicating that low-complexity sequences with high occurrence in host genomes are more likely be misclassified.

**Figure 4.**
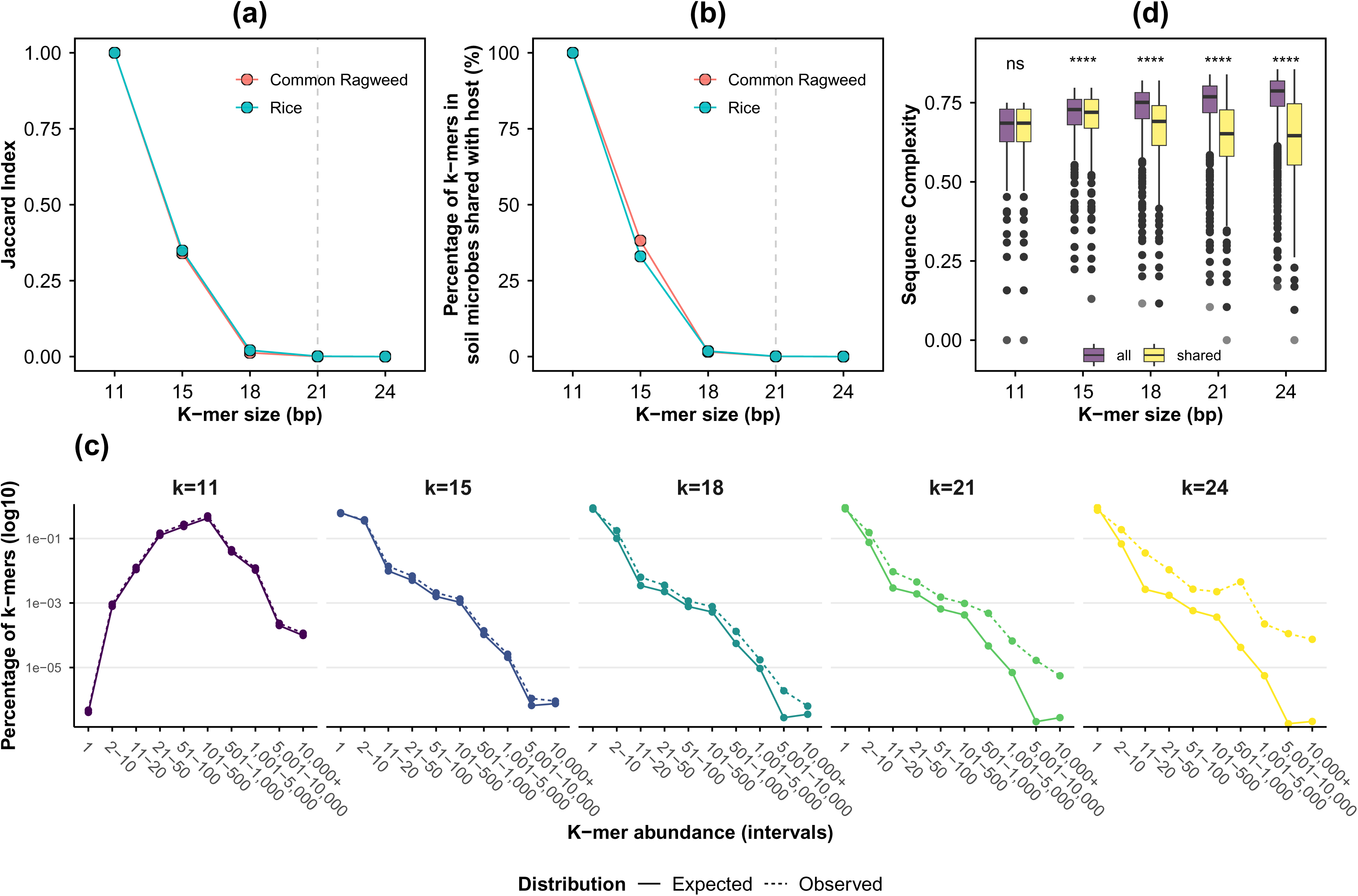
Shared k-mers between host (rice and common ragweed) and representative pan-genomes of the soil microbiome. (a) Jaccard index and (b) the percentage of k-mers shared between host and soil microbiome across different k-mer values using Jellyfish “count” command with the option --mer-len=11, 15, 18, 21 and 24 and with Bloom filter disabled and both strands considered. The decrease of the Jaccard index to 0.02 or below at k=21 (dashed line) was consistent across host species (color-coded lines). (c) The distribution of k-mers in rice (solid line) and those shared between rice and soil microbes (dashed line). K-mers were grouped into intervals (x-axis) based on their occurrence in host genome (1, 2-10, 11-20, etc). (d) Sequence complexity (Wootton-Federhen complexity score) of all (purple) and shared k-mers (yellow) in rice and the soil microbiome across k-mer sizes. *P*-value was obtained from the Wilcoxon signed-rank test.

### Strategies for working with fragmented DNA from historical samples

In addition to host DNA, damage and fragmentation of DNA molecules can affect microbial analyses of historical samples. Because historical samples are scarce and the true microbiome profiles in these samples are unknown, we performed a simulation study to evaluate the effects of host DNA content on metagenomic analyses of the short DNA molecules (20-40 bp) commonly recovered from such samples. A total of 6,000 simulated datasets were generated with *gargammel* for three plant species differing in genomic complexity and size (rice, maize and bread wheat) and 120 representative soil microbes (Supplementary Table 2). To mimic the damage and fragmentation of DNA from historical specimens, the length distribution of one of the herbarium rice samples in this study (Supplementary Figure 7) and nucleotide modifications following the C-to-T substitution dynamics of ancient DNA were incorporated into the simulated sequencing reads. Each dataset contained one million simulated paired-end reads, with host and microbial DNA proportions ranging from 0.01 to 0.45. An average of 0.98 million reads was obtained after quality filtering and merging. The merged read lengths followed a lognormal distribution, ranging from 21 to 150 bp with a peak around 28 bp (Supplementary Figure 7), closely resembling the fragment length profiles typically observed in historical samples. Merged reads were processed with or without host DNA removal, followed by taxonomic classification using custom Kraken2 databases built with k-mer sizes of 21, 24, 28, 31 and 35. Overall, cross-domain misclassification of host reads (assigned to prokaryotes) was very low across plant host species (Figure 5a and Supplementary Figure 8). Without host DNA removal, the highest misclassification rate in rice was about 0.2% at k=21 and k=24, and it dropped to 0.1% or below when host DNA removal was performed. The abundance of soil microbes was consistent between workflows across datasets (Supplementary Figure 9). As expected, the number of classified microbial reads correlated with their proportions (0.01-0.45) specified during read generation, but it was also strongly influenced by the k-mer value of the databases (Figure 5b; Supplementary Figure 10). The highest number of classified microbial reads (about 300,000 for datasets with a bacterial fraction of 0.45) was obtained with databases built with k=24 or k=28, roughly one-third more than that with k=35.

**Figure 5.**
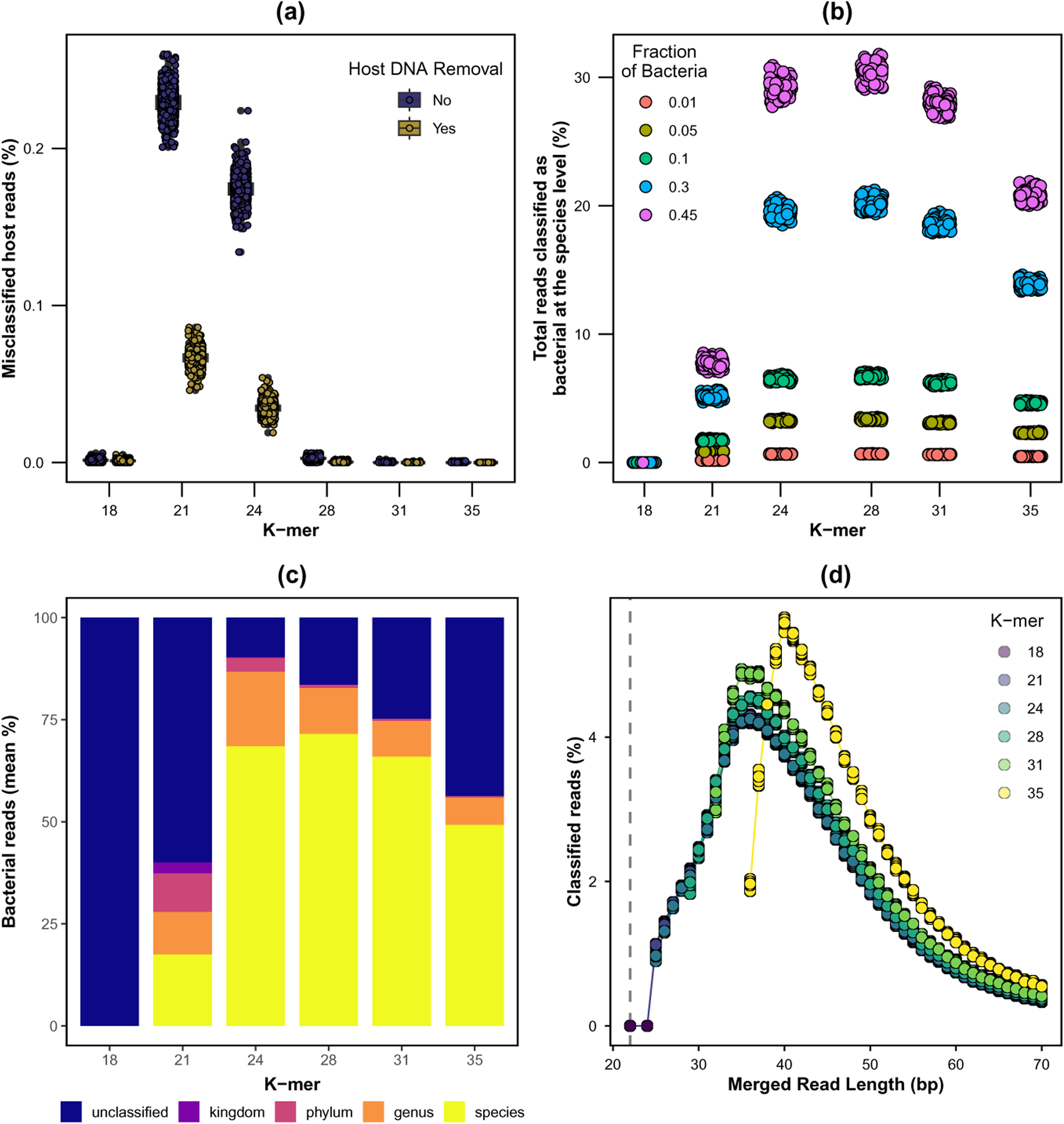
The analyses of the simulated rice datasets annotated with databases of different k-mer values. **(a)** The percentage of host reads misclassified as microbial taxa. The simulated datasets were processed with (gold) and without (purple) host DNA removal step by comparing the mapping results against custom databases built with k=21 to k=35. **(b)** The number of classified bacterial reads in randomly selected 2,000 simulated rice ancient DNA (aDNA) datasets. Merged reads from the simulated datasets were annotated with five databases created with different k-mer values (k= 21, 24, 28, 31 and 35). Subsequent analyses **(c)**-**(d)** were based on the datasets without host DNA removal **(c)** The mean percentage of microbial reads detected in the simulated rice aDNA datasets. Classified reads were color-coded according to their lowest taxonomic rank (species, genus, phylum and kingdom were shown). Unclassified reads were shown in blue. **(d)** The read length distribution of classified reads resolved at the species level across k-mer values.

Reads classified at the species level are generally more informative than those resolved only at higher taxonomic ranks. Among the simulated datasets, the number of microbial reads resolved at the species level ranged from 7,890 to 375,850. The percentage of reads classified at the species level was highest at k=28 (Figure 5c and Supplementary Figure 11). At k=24, a larger percentage of reads was assigned to higher taxonomic ranks (e.g. genus or even phylum), indicating that the databases built with smaller k-mer sizes may improve the overall classification rate, but at the expense of taxonomic resolution.

Because DNA fragments from historical samples are highly variable in length (Supplementary Figures 6 and 12a), we further examined the read length distribution across different k-mer sizes. Regardless of k-mer size, the length distribution of classified microbial reads was lognormal with a peak length at 30-40 bp (Figure 5d), following the read length distribution of the herbarium rice sample used to generate the simulation datasets. The shortest reads that could be annotated were constrained by the k-mer size used to build the database (e.g. the lower bound of read length for k=35 is 35 bp), because Kraken2 cannot classify reads shorter than the k-mer size. These results suggest that annotation with two or more k-mer sizes could extend the coverage across the wider spectrum of fragment lengths commonly found in historical specimens.

### Improving recall rate of microbial signals with two-step classification

To determine the optimal annotation strategy, we compared the number of reads classified at the species level across different combinations of k-mer sizes in the simulated datasets. In the single-step workflow (Figure 1), reads were annotated with a custom Kraken2 database (Standard databases built with k=21, 24, 28, 31 and 35). With a k=35 database, only 47.7% of microbial reads were classified at the species level (Figure 6b-c). Publicly available Kraken2 databases (e.g. PlusPFP) are built with a k=35, suggesting that they may not be optimal for annotating reads from historical or ancient samples. When a k=28 database was used instead, the average classification rate increased to 67.1%. However, using a smaller k-mer size may inflate cross-domain misclassification, especially when host DNA is not removed (Figure 5a). To address this, we propose a two-step annotation workflow (Figure 6a) in which reads are first annotated with a database built with a large k-mer size (e.g. 31 or 35). Reads that are unclassified or resolved only at the genus level or above in the first step are then processed against a second database built with a smaller k-mer size (e.g. 28 or 24). This strategy achieved a higher classification rate (71.7%), outperforming the single-step workflow. As expected, recall rate was strongly governed by the choice of k-mer size(s): it ranged from 0.04 to 0.66 In the single-step workflow, compared with 0.49-0.74 in the two-step workflow (Figure 6c). In contrast, the precision of taxonomic classification remained consistent between 0.97 and 0.99, regardless of k-mer size (Figure 6c and Supplementary Table 7). When the herbarium rice samples were analysed with the two-step approach, more reads were classified than with the single-step approach (Supplementary Figure 12b).

**Figure 6.**
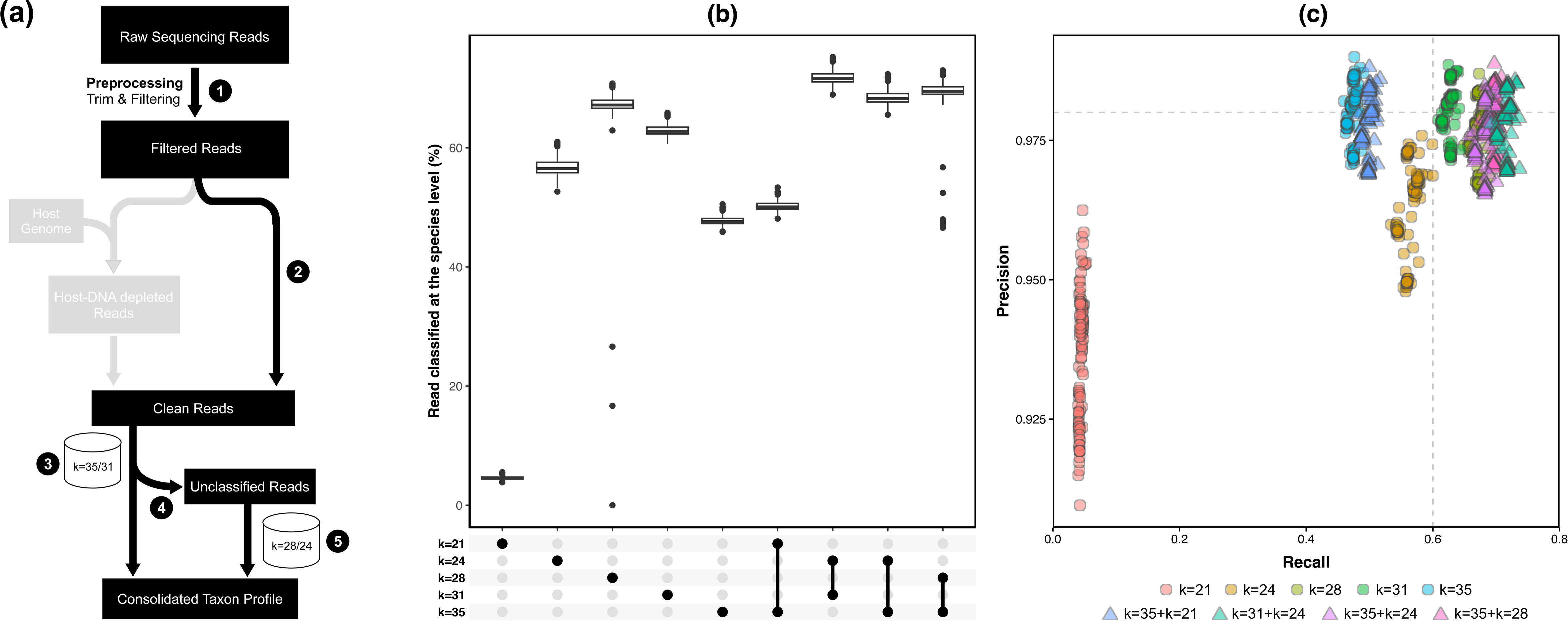
**(a)** Proposed two-step analysis workflow for the taxonomic annotation of metagenomic reads in historical samples without host DNA removal. 1. raw sequencing data preprocessing: trimming or quality control filtering; 2. Other preprocessing steps of the sequencing reads such as merging of paired-end reads or removal of low-complexity or repetitive sequences; 3. The first stage of classification with an annotation database built with a large k-mer value (k=35 or k=31); 4. Unclassified reads or those assigned with a non-species level taxonomy ID can be processed with the second stage of classification; 5. Classification of unclassified reads with an annotated database built with a shorter k-mer value (k28/k24). Subsequent analyses **(b)**-**(c)** were performed on the simulated rice datasets without host DNA removal. **(b)** The percentage of microbial reads identified by single-or two-step search strategies in the simulated rice datasets. The k-mer value(s) used for the annotation is highlighted with a black circle in the UpSet plot below. **(c)** The precision and recall of 100 simulated rice datasets (randomly selected out of 2,000 datasets) analysed in single and two-step analysis. Points were color-coded based on the combinations of k-mer value(s) (k=21, k=24, k=28, k=31, k=35, k=31+k=24, k=35+k=21, k=35+k=24 and k=35+k=28) used. Data processed with single-step analysis workflow are shown as circles while those from two-step workflow as triangles. The precision and recall for all 2,000 simulated rice datasets were included in Supplementary Table 7.

## Discussion

Historical specimens are valuable for understanding how host-associated microbial communities have changed over time and in response to environmental changes. However, profiling host-associated microbiomes in historical samples is challenging due to contaminating DNA from host species or other non-target organisms. These extraneous sequences have been suggested to interfere with downstream analyses, potentially leading to false identification of microbial signals or misassembly of sequence contigs (Schmieder & Edwards, 2011). Host DNA removal becomes infeasible when reference genomes are unavailable, as is often the case for non-model species or extinct organisms. This raises the question of the extent to which microbiome analyses of historical specimens remain informative in the absence of host DNA depletion. To address this, alongside simulation and k-mer analyses, we analysed a large cohort of metagenomes spanning historical and contemporary specimens from both model and non-model organisms. Our findings demonstrate that host DNA has little to no effect on microbial analyses, regardless of its abundance.

Without removing host DNA, sequences from host genomes may be falsely classified as microbial. The findings from the k-mer analysis and the simulation study suggest that false positives due to misclassification of host DNA could only become observable when metagenomic reads were annotated with databases built with a small k-mer value (e.g. k=24 or smaller). Nonetheless, the highest false positive rate (∼0.25% of host reads at k=21) was observed in rice while extremely low values (0.006-0.075%) were detected for maize and bread wheat. In sum, host DNA removal may not be as critical as previously believed and misclassification could be reduced by downstream bioinformatics workflows such as the two-step classification workflow proposed in this study.

DNA recovered from historical specimens or archaeological artifacts is often degraded and highly fragmented (Burbano & Gutaker, 2023). Damage patterns due to chemical lesions (e.g. deamination) could be repaired experimentally prior to library preparation and sequencing while there is no existing method to reverse fragmentation. Fragmented DNA molecules complicate the library preparation and downstream bioinformatic analyses. The typical workflow for profiling metagenomic reads in historical specimens relies on k-mer-based classification tools such as Kraken2 and Centrifuge (Kim et al., 2016). The sensitivity and specificity of these tools can be adjusted through k-mer size or seed length. The publicly available Kraken2 databases were built with a k-mer value of 35 (Arizmendi Cardenas et al., 2022; Lu et al., 2022). This setting is ideal for general use as it optimises specificity and sensitivity but is not optimal for short DNA sequences as confirmed by our results, in which the recall rate at k=28 was notably higher than that at k=35. Several k-mer values have been proposed for various scenarios and types of datasets. A k-mer size of 29 was suggested to build databases for filtering exogenous DNA in historical samples (Ravishankar et al., 2024). For metagenomic datasets, a k-mer size of 21 was shown to be sufficient to obtain accurate results in most cases (Ondov et al., 2016).

Nevertheless, annotation using a single database with a fixed, short k-mer value may be suitable only for limited use-case scenarios, as fragmented DNA molecules from historical specimens span a wide range of lengths, from 20 to over 100 bp. To cope with this, a two-step annotation workflow was proposed in this study to improve the recall rate of short DNA fragments. The simulation study and results of applying the two-step workflow (e.g. k=31/35 and k=28) to herbarium rice samples showed that a notable improvement in recall rate was achieved without sacrificing the precision. The optimal combination of k-mer values should be determined on a case-by-case basis based on the length profile of DNA molecules recovered from the historical samples in question. This improvement would enable the recovery of more informative signals from precious historical samples that might otherwise be discarded.

Beyond host DNA removal and fragmented DNA molecules, which were the primary focus of this study, several other aspects merit consideration when analysing metagenomic datasets from historical specimens. First, as with metagenomes from other sample types (e.g. clinical specimens), taxon filtering can have a substantial impact on downstream microbiome data analysis and on reduction of technical variability. Proper taxon filtering can reduce the number of falsely identified taxa, which are typically present at low abundance and prevalence (Cao et al., 2020). Second, shared k-mers between host and microbes were of low complexity, suggesting that repetitive elements or other low-complexity motifs in eukaryotic genomes may contribute to false positives in taxonomic classification. A sequence complexity filter or masking of repetitive elements prior to taxonomic classification may help mitigate this issue. Third, host DNA depletion could reduce the computational burden and complexity when functional profiling of metagenomic reads or *de novo* assembly of novel microbial genomes is the prime objective. In such cases, genomes of closely or distantly related species may aid in removing host-derived reads. As such, a more lenient mapping strategy (e.g. reducing the seed length or lowering mismatch and gap penalty) may be beneficial. Fourth, assessing the mapping evenness of classified reads could help identify false positives that evade taxon filtering steps. Other technical factors, such as sample size and sequencing depth, can also influence the identification of differential taxa (Nearing et al., 2022). However, we did not examine these factors in this study, as they are often beyond investigators’ control when studying historical samples.

In summary, informative microbial analyses could still be performed on historical samples even when genomic sequences of host species are unavailable. This approach may be applicable to exploring the microbial ecology of non-model organisms. We also showed that short microbial sequences (as short as 24 bp) can be distinguished from those derived from the host genome and therefore informative results can still be obtained. To cope with short DNA fragments, a two-step annotation workflow (e.g. reads annotated with a k=31 database followed by a k=28/24 database) was proposed. Although our findings suggest that the genome size and complexity may not critically affect downstream microbiome analyses, a comprehensive evaluation of the analytic workflow remains warranted. Finally, contamination arising from post-mortem colonization can accumulate over time. In this study, we did not separate the contamination from the reads derived from host-associated microbes. Our findings were not critically affected by the sources of the microbes (either host-associated or contamination) as host DNA content and optimal analysis strategies on microbial signals were the prime focus of this study. Future work should focus on the detection of microbial contaminants in historical samples through various approaches such as DNA damage patterns (especially for recent contamination) or a catalogue of post-mortem contaminants commonly detected in museum collections. Other aspects worth exploring, yet overlooked in this study, are the availability of microbial reference genomes in the annotation database and the use of host genomes that may contain contaminant microbial sequences. Historical samples may harbor microbial taxa that have become rare due to anthropogenic pressures or are entirely absent from contemporary samples. How to effectively detect and characterize these poorly represented or unknown microbial taxa remains an open question. Overall, this study marks an important milestone in enabling metagenomic investigation of historical samples, contributing to our understanding of how anthropogenic changes have shaped hosts and their associated microbiomes.

## Data Availability Statement

Raw sequence reads generated in this study are deposited in EBI ENA under the accession PRJEB87646.

## Acknowledgements

SKN and RMG are both supported by UKRI Frontier Research Guarantee grant EP/X022404/1 to RMG. The authors acknowledge Research Computing at the James Hutton Institute for providing computational resources and technical support for the “UK’s Crop Diversity Bioinformatics HPC” (BBSRC grants BB/S019669/1 and BB/X019683/1), use of which has contributed to the results reported within this paper (Percival-Alwyn et al., 2025). The authors also acknowledge help from Robin Becina Guevarra with early versions of the manuscript.

## Ethical Statement

This study did not involve any human or animal subjects. Our work follows research integrity standards.

## Non-monetary Benefit-sharing Statement

Collaboration, cooperation and contribution in scientific research and development programmes, particularly biotechnological research activities, where possible in the Party providing genetic resources.

## Author Contributions

RMG conceptualized the study with inputs from SKN. SKN conceptualized the k-mer comparison approach. SKN carried out analyses and designed the figures with inputs from RMG. SKN wrote the manuscript with inputs from RMG.

