## Supplementary material for "Effects of host DNA and k-mer size on metagenomic classification of historical samples": Table 1

| **Dataset** | **Sample Size (N)** ^§^ | **With Host DNA Removal** | | |  | **Without Host Removal** | | | **Taxa Removed by Host DNA Removal ^#^** | **Taxon# with ≥10% Change in Mean Read Count between Workflows** |  | **Mean Read Count** | |
| --- | --- | --- | --- | --- | --- | --- | --- | --- | --- | --- | --- | --- | --- |
|  |  | Before filtering **^¶^** | After filtering | % filtered |  | Before filtering | After filtering | % filtered |  |  |  | taxon with ≥10% change | taxon with <10% change |
| Contemporary Rice | 330 | 4,460 | 390 | 91.3% |  | 4,614 | 390 | 91.6% | 0 (0.0%) | 55 (14.1%) |  | 1,951.6 | 18,157.6 |
| Herbarium Rice | 95 | 8,057 | 853 | 89.4% |  | 8,731 | 855 | 90.2% | 2 (0.3%) | 7 (0.8%) |  | 90.6 | 467.0 |
| Herbarium Ragweed | 37 | 4,049 | 184 | 95.5% |  | 4,315 | 185 | 95.7% | 1 (0.5%) | 0 (0.0%) |  | - | 616.2 |
| Museum Bumblebee | 402 | 3,978 | 113 | 97.2% |  | 4,497 | 113 | 97.0% | 0 (0.0%) | 0 (0.0%) |  | - | 12,543.8 |

^§^ Sample Size (N): no sample failed in host DNA removal step

**^¶^** Filtering cutoff: read count = 50 (100 for contemporary rice dataset), prevalence = 2% of samples

**^#^** taxon removed: the number of taxa detected between the workflow with and without host DNA removal step before filtering
