## Supplementary Figures for "Effects of host DNA and k-mer size on metagenomic classification of historical samples"

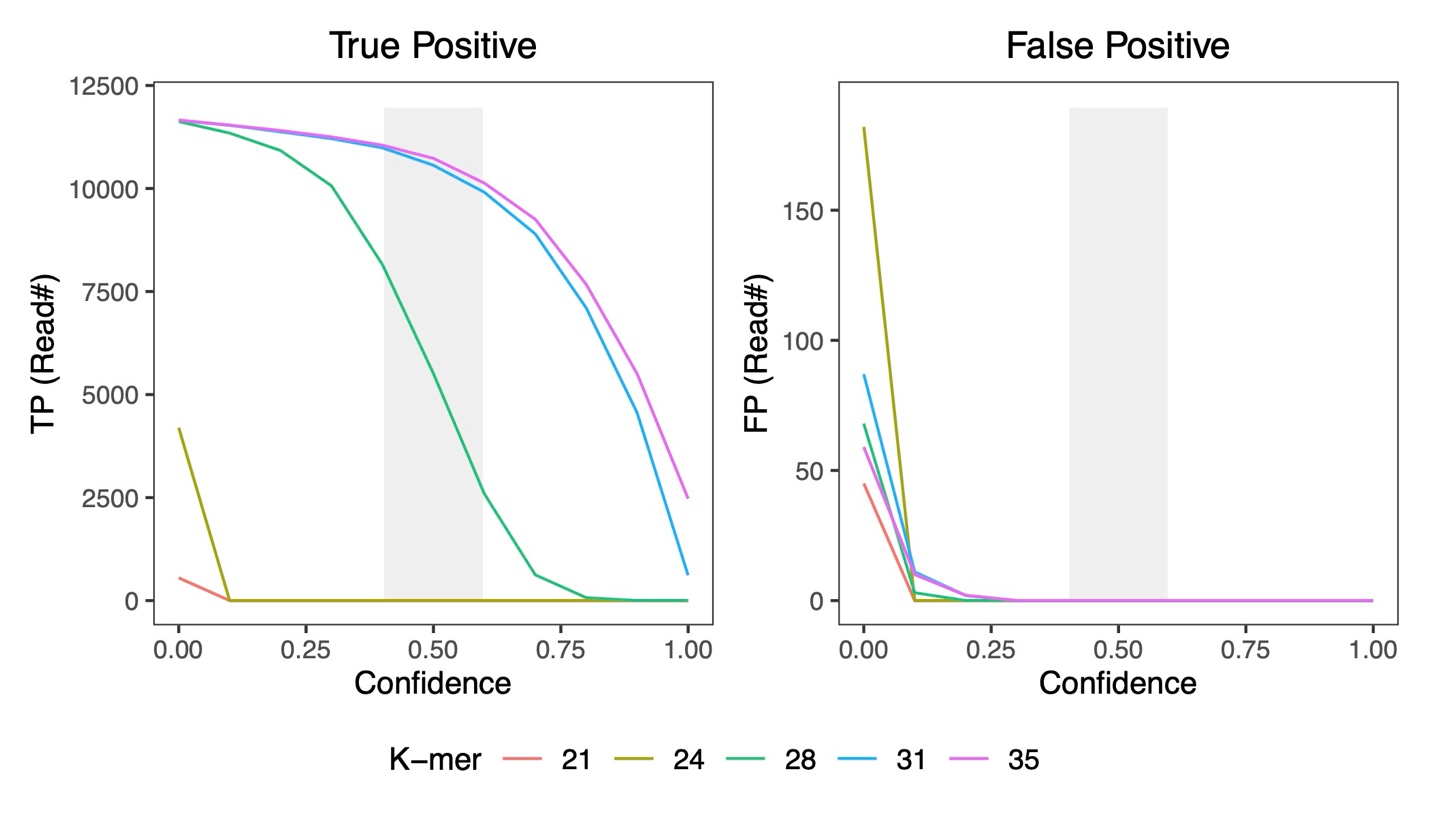

Supplementary Figure 1. Estimation of the optimal confidence values for taxonomic classification with Kraken2. A simulated data of 12,500 pair-end reads with a length of 100bp was generated from the reference genomes of *Staphylococcus epidermidis* strain ATCC 14900 (NZ_CP035288.1) using ART_illumina v2.5.8. Five standard-16 databases were built with k-mer values of 21, 24, 28, 31, and 35 (color-coded). The simulated reads were mapped against the databases using Kraken2 with confidence values of 0.0-1.0 at intervals of 0.1. Reads classified to *S. epidermidis* were true positive (TP) while those mapped to species other than *S. epidermidis* were considered as false positive (FP). The optimal range of confidence values (0.4-0.6) is highlighted with a grey shade.

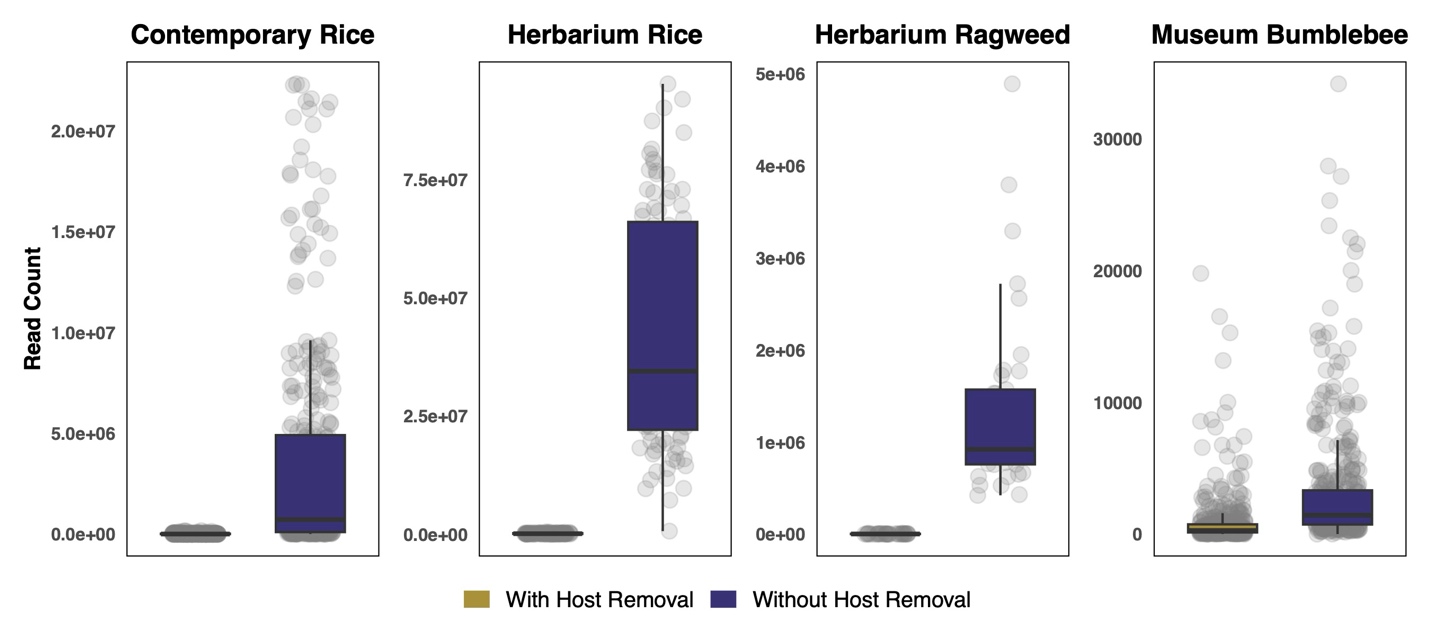

Supplementary Figure 2. Proportion of host DNA content in each sample (point in grey) across datasets between the workflows with (gold) or without (purple) host DNA removal using bowtie2. Some residual host DNA was detected in the downstream analysis with Kraken2.

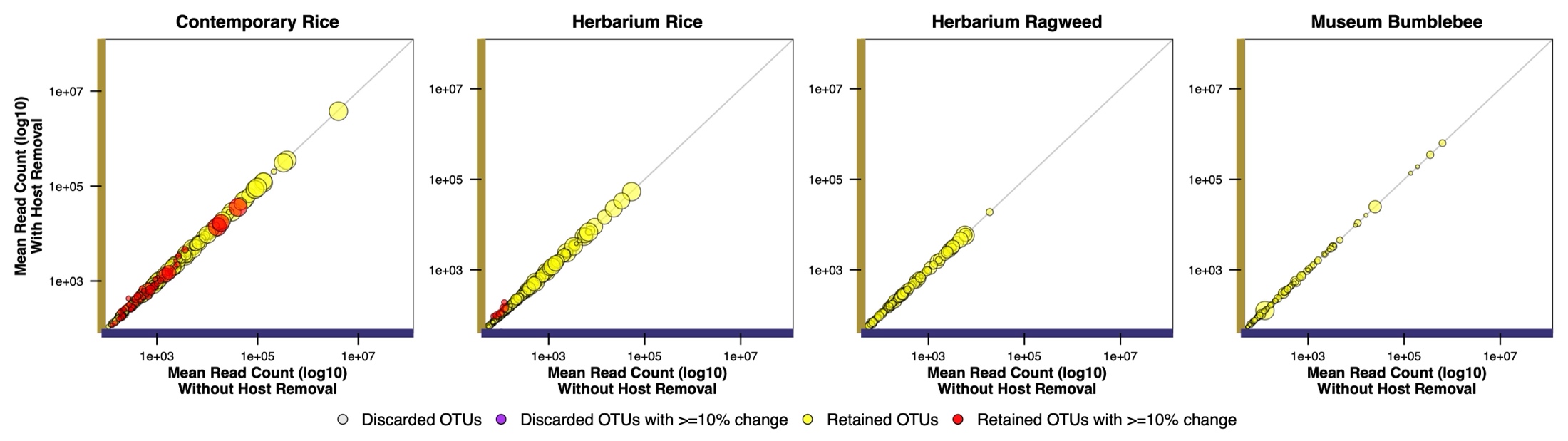

Supplementary Figure 3. The distribution of bacterial taxon in assigned read count (log10) between two workflows with (y-axis; gold) and without (x-axis; purple) host DNA removal. Rare taxa (highlighted in grey color) were excluded in the downstream analyses. Each circle represents a bacterial taxon. Those with more than 10% change in read count between two workflows are highlighted in red. The size of the circle represents the prevalence of the taxa.

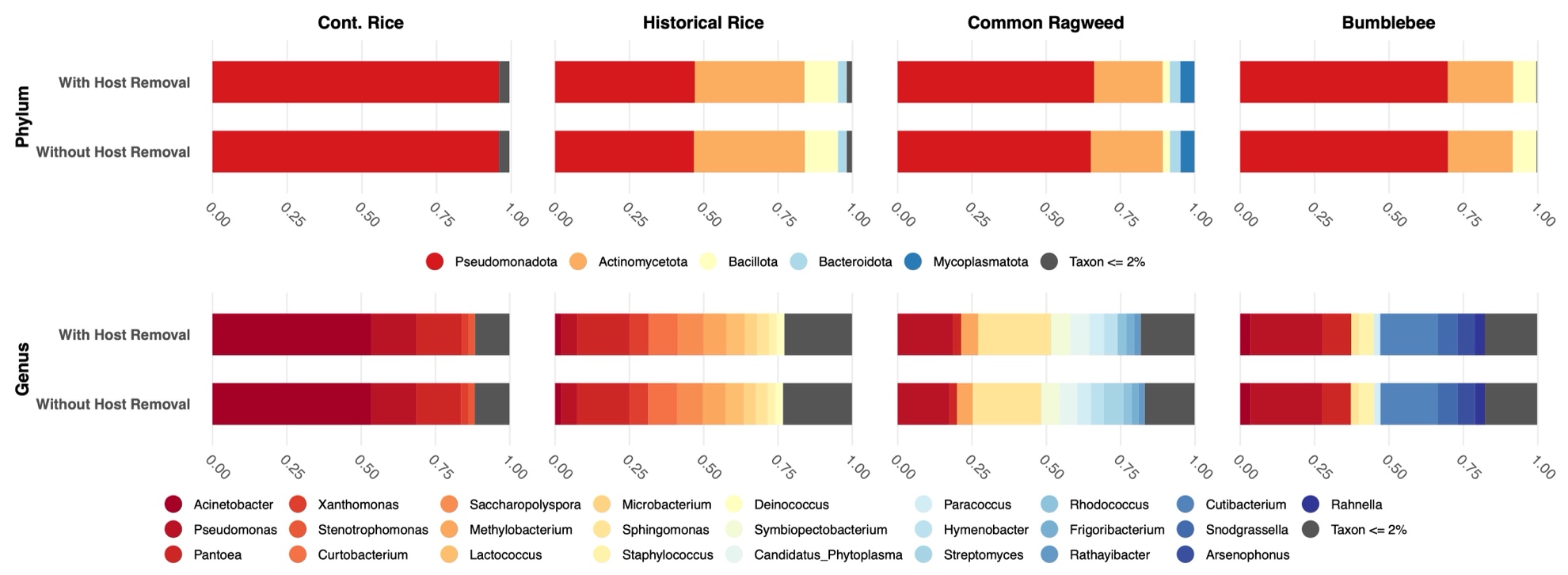

Supplementary Figure 4. Stacked-bar charts showing the relative abundance of top bacterial taxa at different taxonomic ranks between the two workflows (with or without host DNA removal) at the Phylum (up) and Genus (bottom) level. Taxa (grey) with an abundance less than the thresholds were grouped together.

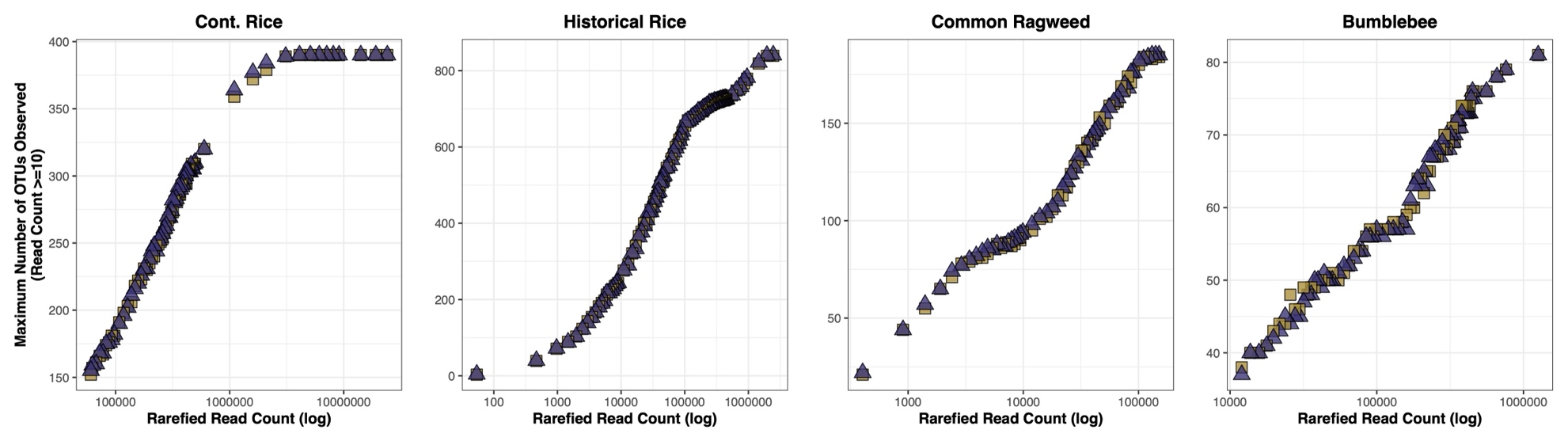

Supplementary Figure 5. The maximum number of detected taxa for each dataset in rarefaction analyses between two workflows with (square in gold color) and without (triangle in purple color) host DNA removal.

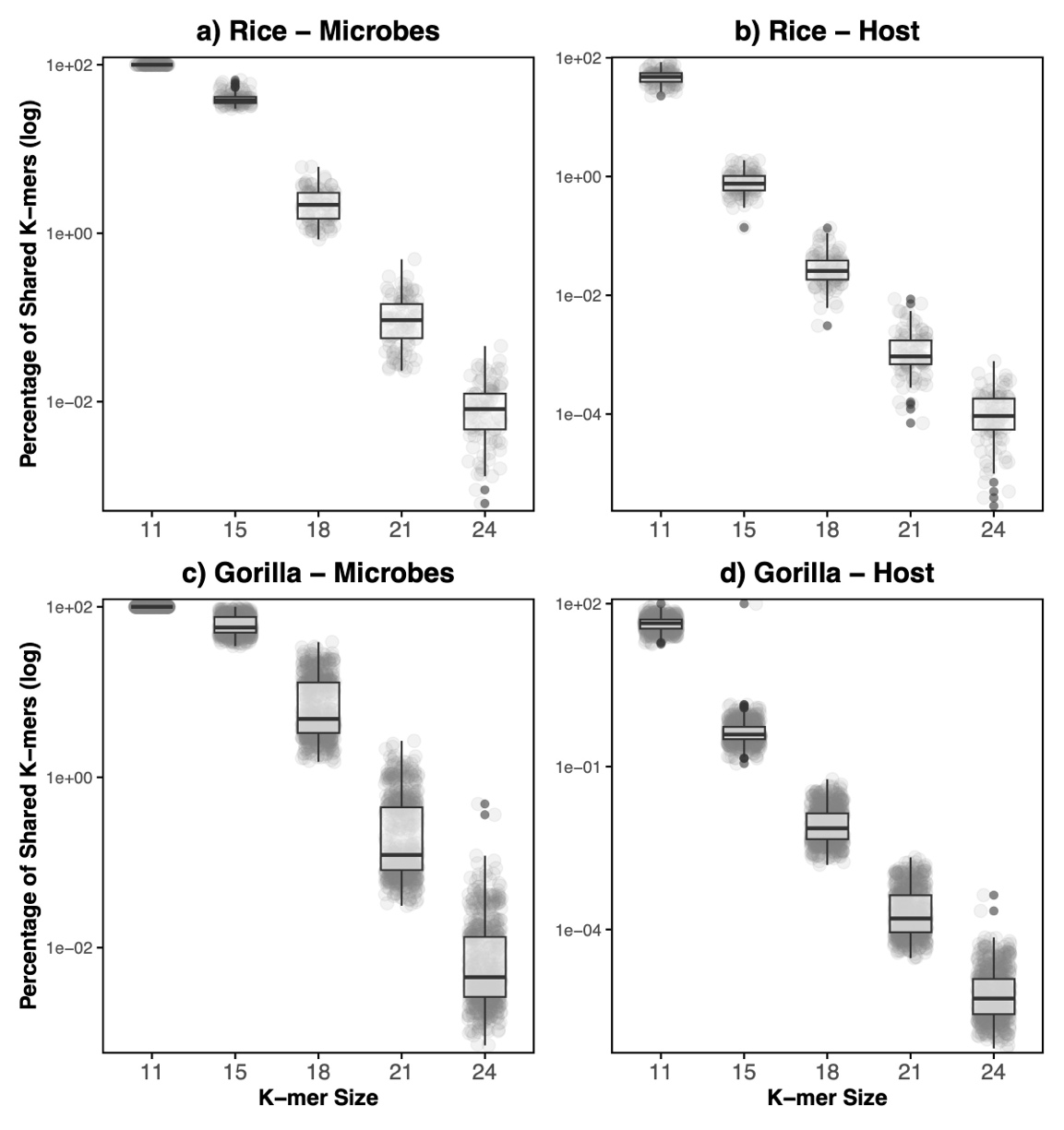

Supplementary Figure 6. Log-transformed percentage of shared k-mers across k-mer size (k=11, 15, 18, 21 and 24) in (a) soil microbiota, (b) rice.

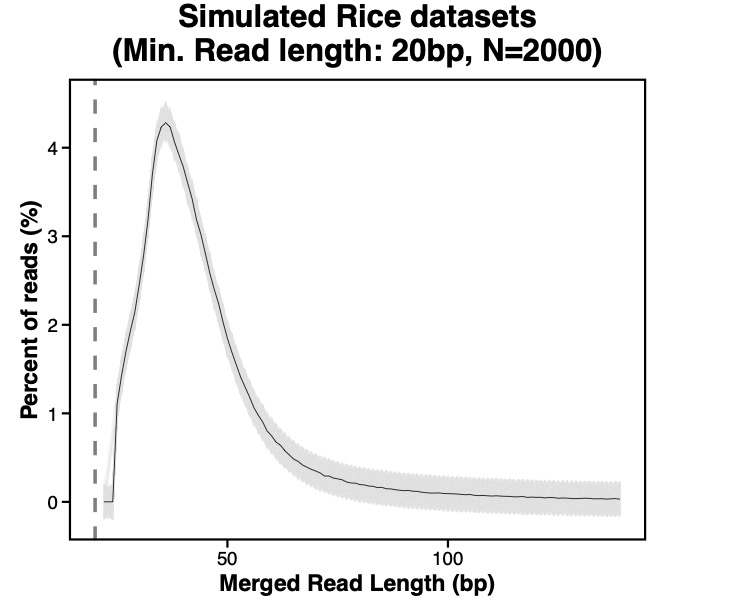

Supplementary Figure 7. The read length distribution of 1-million merged reads summarized from 2,000 simulated aDNA rice datasets. The average was shown as a solid line.

| (a) | **(b)** |
| --- | --- |
| 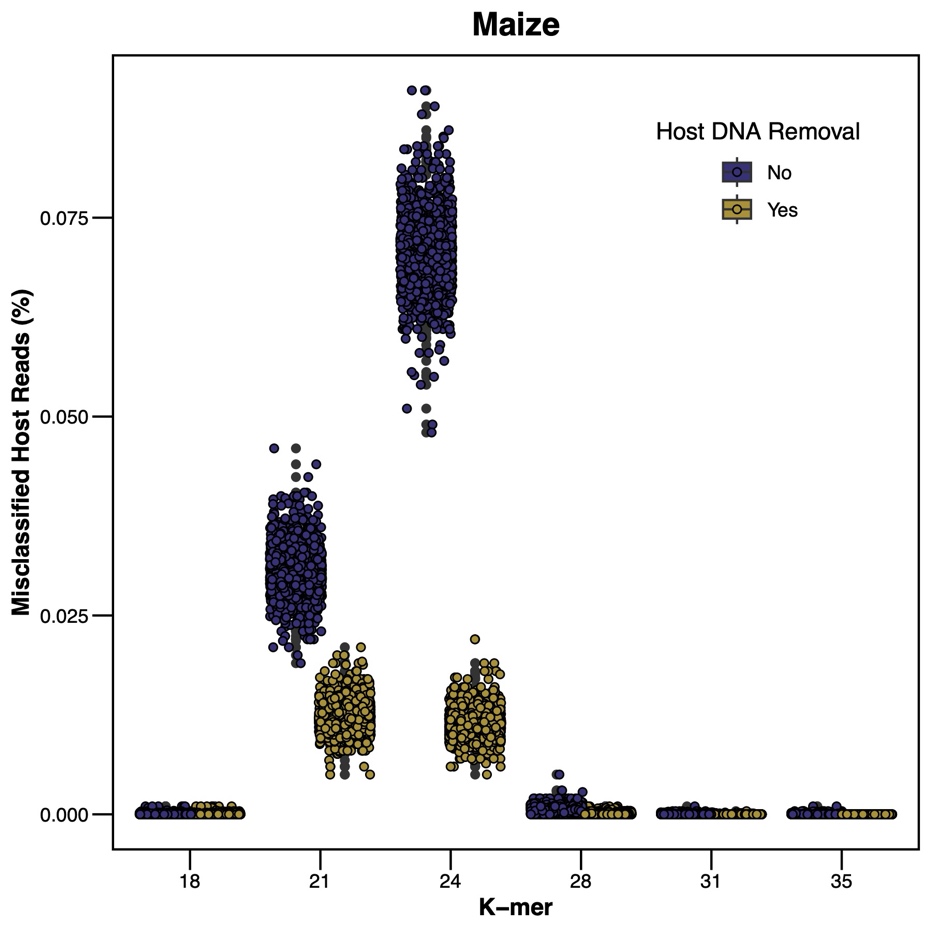 | 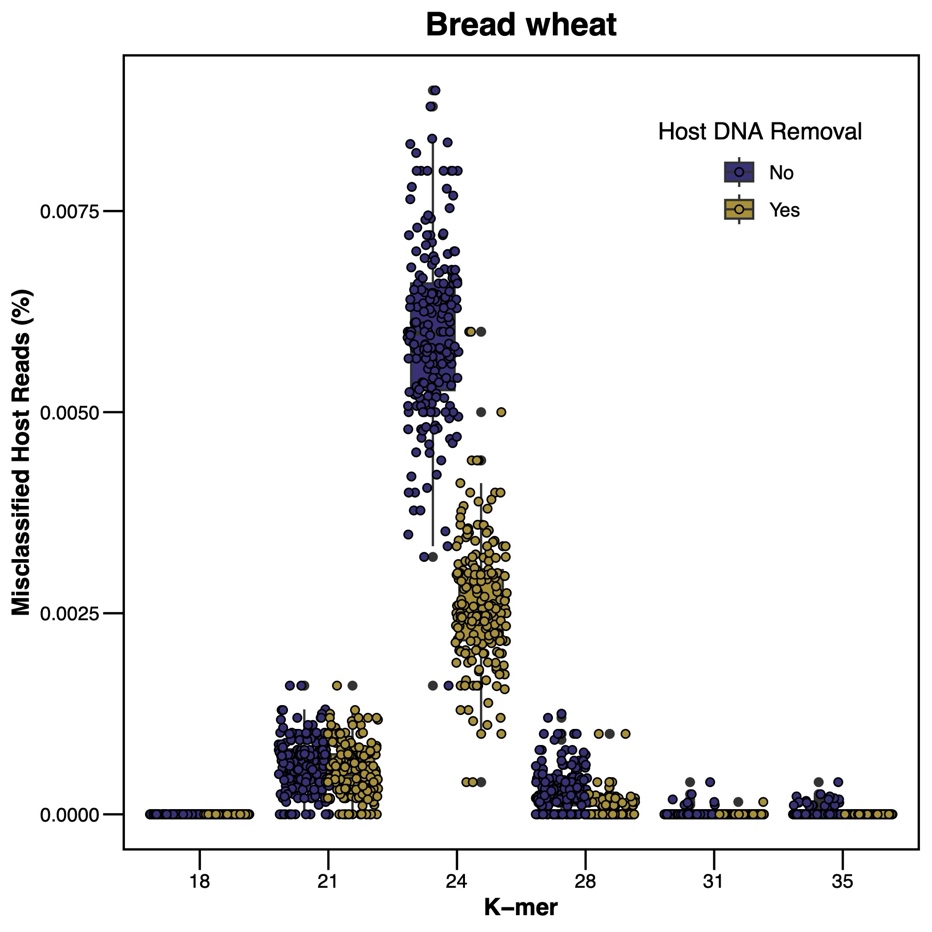 |

Supplementary Figure 8. The percentage of host reads misclassified as microbial taxa in the simulated **(A)** maize and **(B)** bread wheat datasets. Merged reads from 2,000 simulated datasets were annotated with databases created with different k-mer sizes (18, 21, 24, 28, 31 and 35). The simulated datasets were processed with (gold) and without (purple) host DNA content removal step.

**(A)**

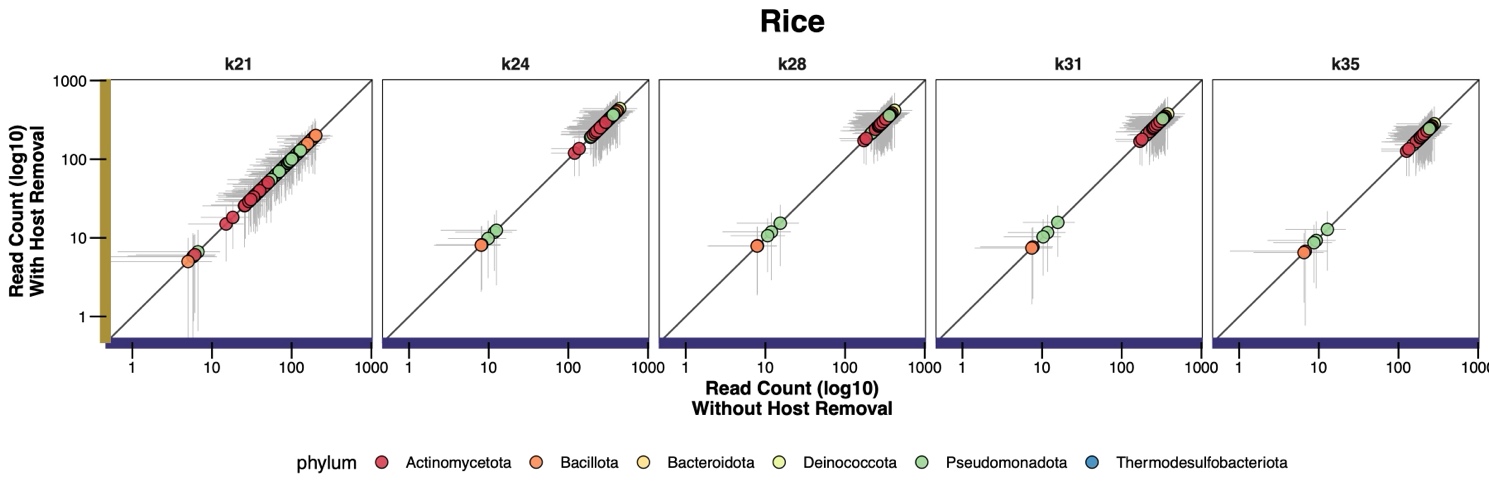

**(B)**

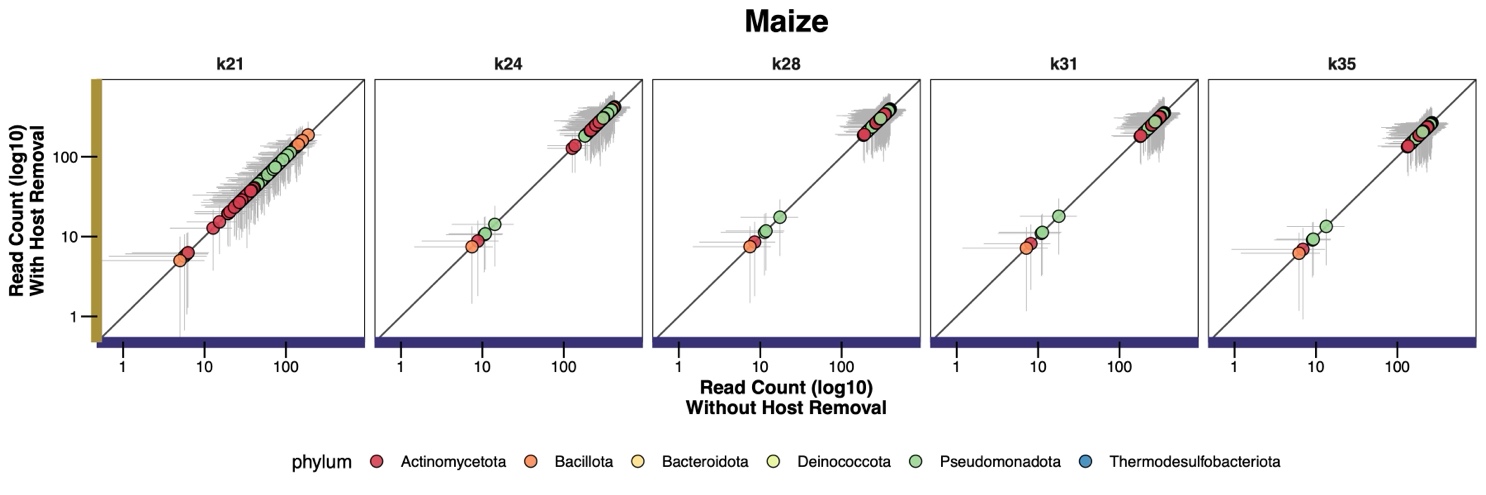

**(C)**

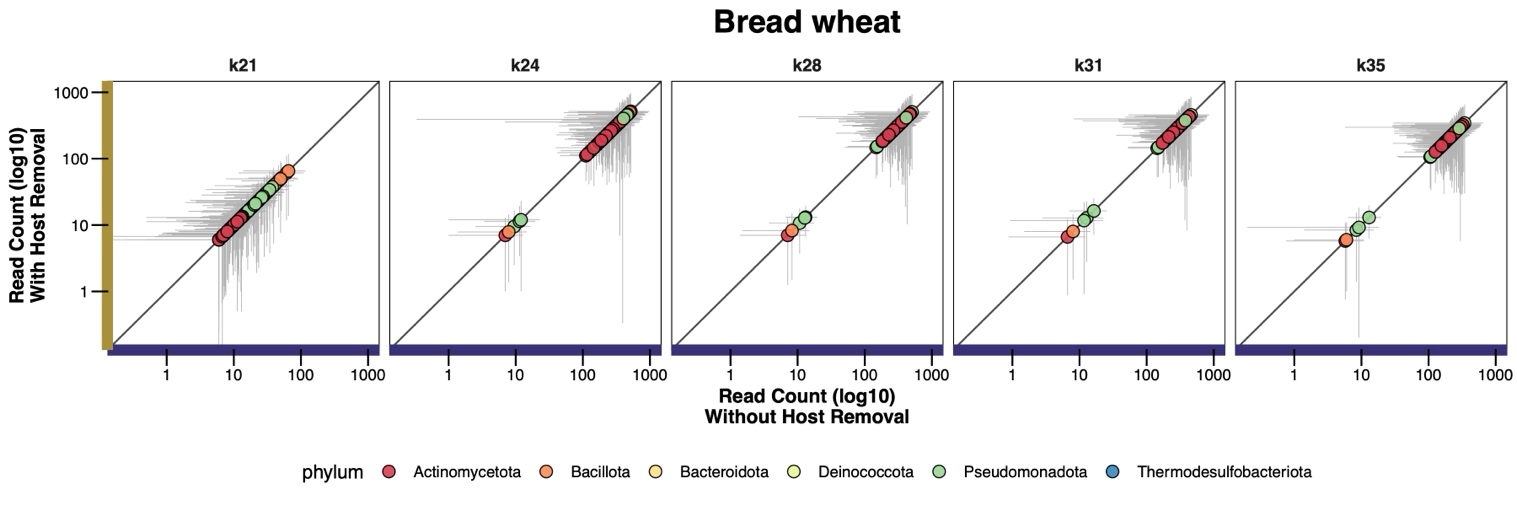

Supplementary Figure 9. The distribution of soil microbes between two workflows with (y-axis; log-transformed; gold) and without (x-axis; log-transformed; purple) host DNA removal in **(A)** simulated rice datasets, **(B)** simulated maize datasets, and **(C)** simulated wheat datasets. Merged reads from each dataset was annotated with databases built with different k-mer values (from k=21 to k=35). K=18 was not included as there was no read assigned to the species level. Each circle represented one of 107 soil microbes summarized from 100 simulated aDNA datasets and was color-coded by their phylum. The vertical and horizontal grey lines behind each circle indicated the first quantile of the read counts summarized from the simulated datasets.

(A)

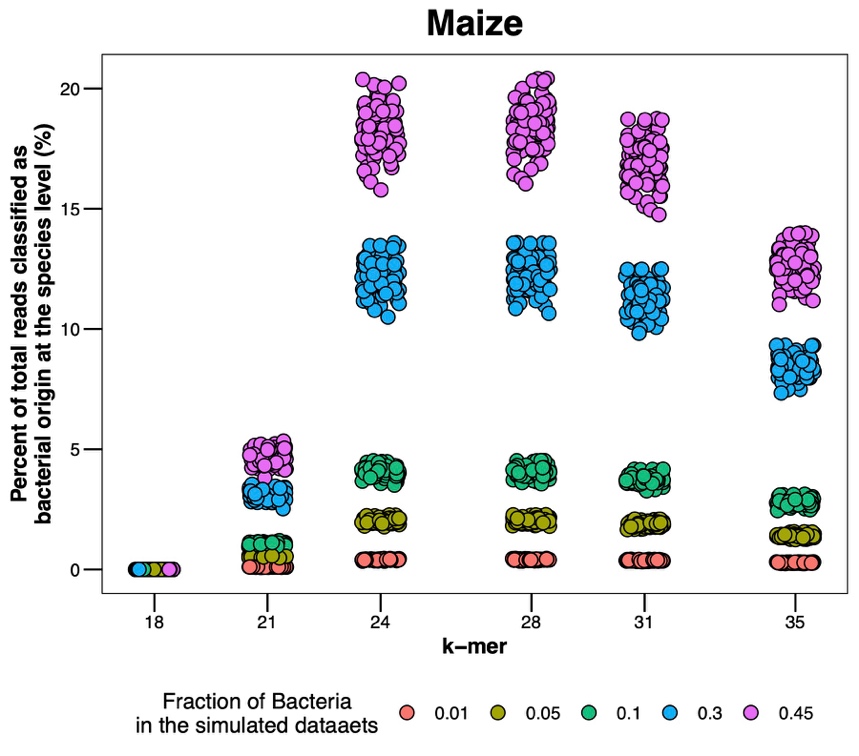

(B)

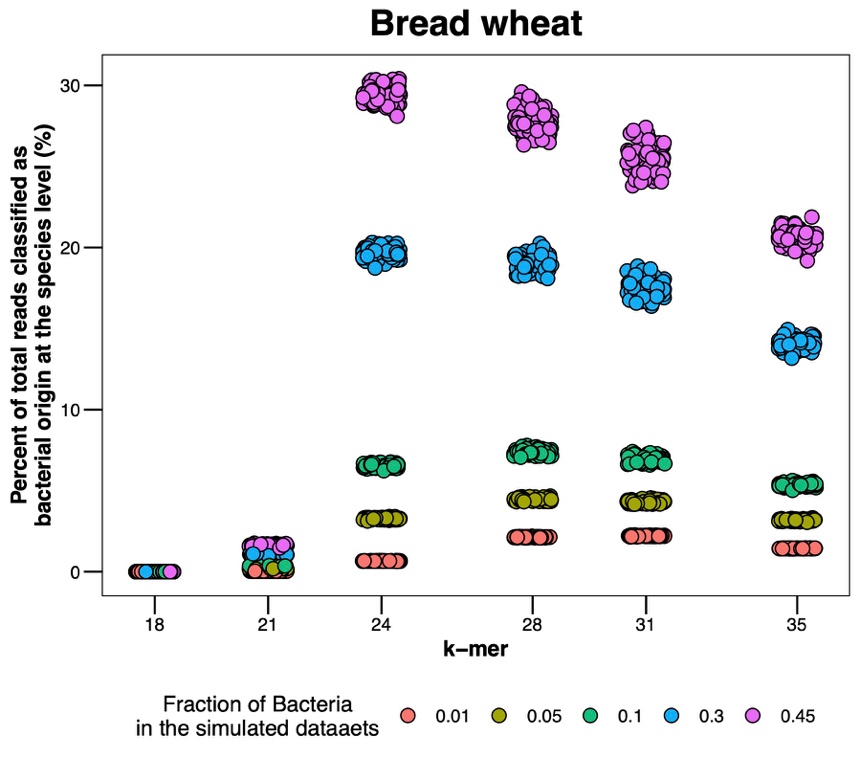

Supplementary Figure 10. The number of classified reads in simulated (A) maize and (B) bread wheat aDNA datasets. The fraction of reads assigned to bacteria is color-coded.

(A)

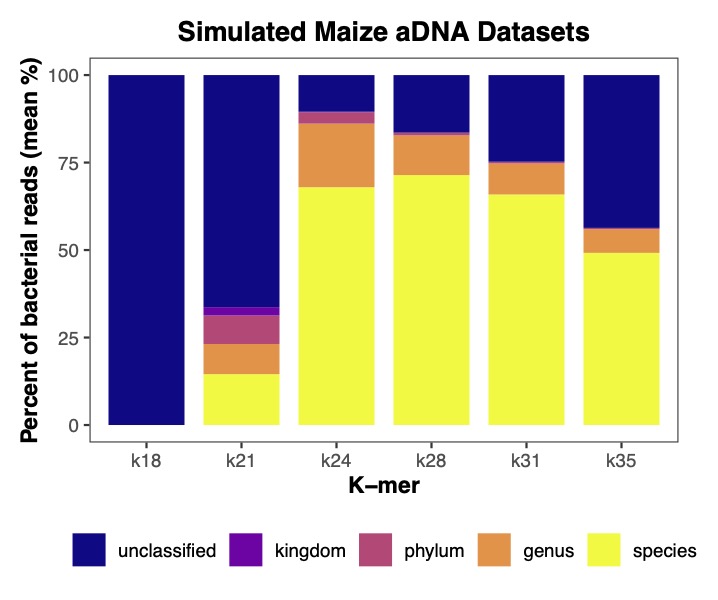

(B)

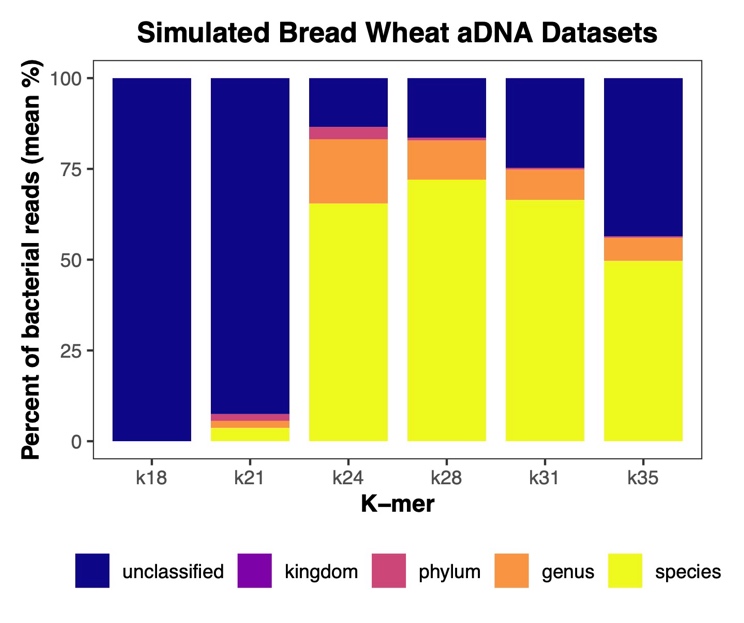

Supplementary Figure 11. The percentage of classified reads assigned at the lowest taxonomic ranks in the simulated (A) maize and (B) bread wheat aDNA datasets.

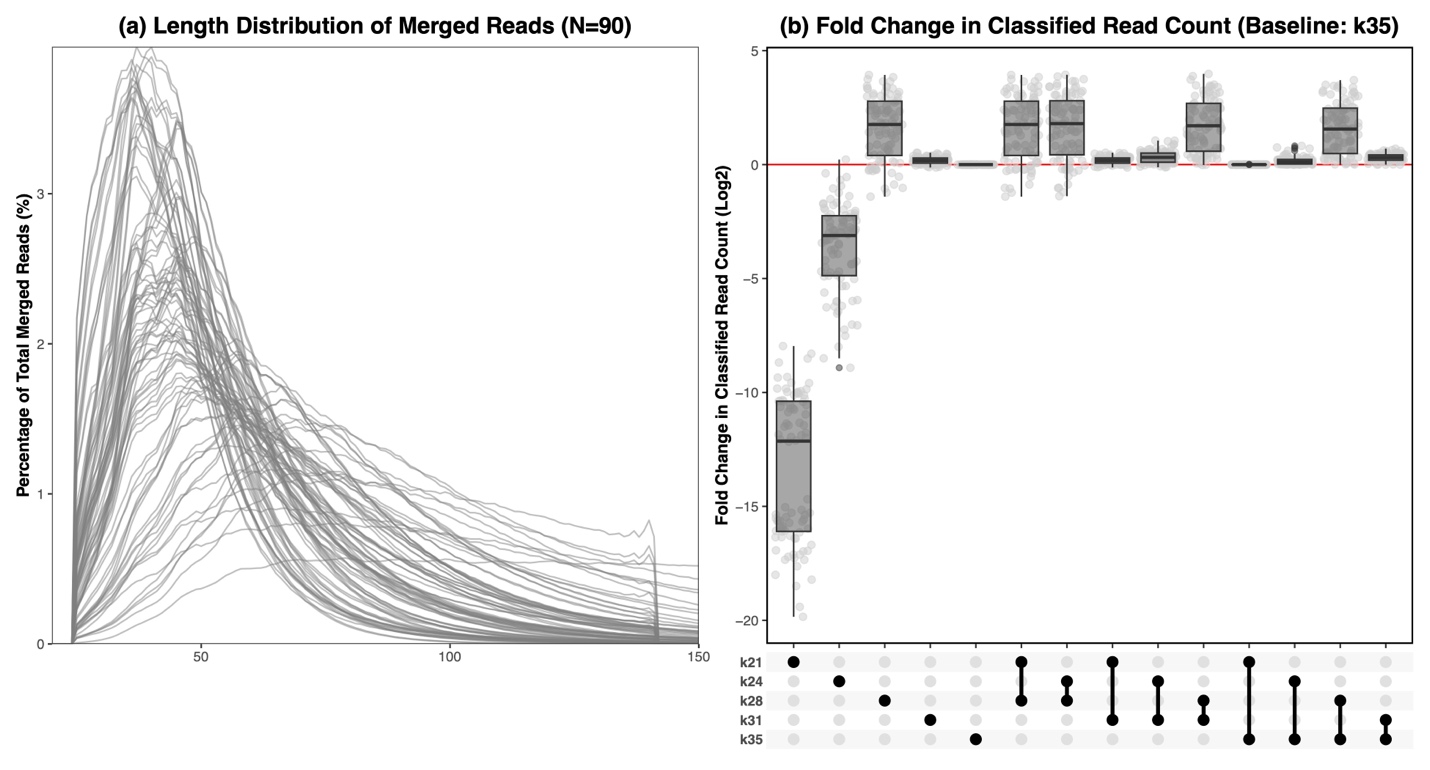

Supplementary Figure 12. Comparison of single and two-step classification for herbarium rice samples (N=90). **(a)** The length distribution of merged reads in percentage **(b)** The fold change in the number of classified reads resolved at species level for the herbarium rice samples. The number of classified reads annotated with a k=35 Standard-16 database was taken as a baseline (red line). The log2 fold changes were calculated as the ratio of reads classified by the single or two-step method to their baseline value, followed by log2 transformation.
